# Localization of signaling receptors maximizes cellular information acquisition in spatially-structured natural environments

**DOI:** 10.1101/2021.07.01.450787

**Authors:** Zitong Jerry Wang, Matt Thomson

**Affiliations:** Division of Biology and Biological Engineering, California Institute of Technology, Pasadena, California, 91125, USA

**Keywords:** spatial organization, natural environmental statistics, cell navigation, information processing, cell sensing

## Abstract

Cells in natural environments like tissue or soil sense and respond to extracellular ligands with intricately structured and non-monotonic spatial distributions, sculpted by processes such as fluid flow and substrate adhesion. In this work, we show that spatial sensing and navigation can be optimized by adapting the spatial organization of signaling pathways to the spatial structure of the environment. We develop an information-theoretic framework for computing the optimal spatial organization of a sensing system for a given signaling environment. We find that receptor localization maximizes information acquisition in simulated natural contexts, including tissue and soil. Receptor localization extends naturally to produce a dynamic protocol for continuously redistributing signaling receptors, which when implemented using simple feedback, boosts cell navigation efficiency by 30-fold. Broadly, our work shows how cells can maximize the fidelity of information transfer by adapting the spatial organization of signaling molecules to the spatial structure of the environment.

## Introduction

Cells sense and respond in spatially-structured environments, where signal distributions are determined by various chemical and physical processes such as substrate binding and fluid flow [1]. In tissue and soil, distributions of extracellular ligands can be spatially discontinuous, consisting of local ligand patches [2–14]. In tissue, diffusive signaling molecules are transported by interstitial fluid through a porous medium. These molecules are then captured by cells and a non-uniform network of extracellular matrix (ECM) fibers, taking on a stable and highly reticulated distribution [2–4, 6–8]. For example, ECM-bound chemokine (CCL21) gradients extending from lymphatic vessels take on stable spatial structures, characterized by regions of high ligand concentration separated by spatial discontinuities [3]. Similar observations have been made for the distribution of other chemokines, axon guidance cues, and morphogens in tissues [6–8, 13]. In soil, a heterogeneous pore network influences the spatial distribution of nutrients by dictating both the locations of nutrient sources as well as where nutrients likely accumulate [9–12]. Free-living cells detect chemical cues released by patchy distributions of microorganisms, where molecules are moved via fluid flow and diffusion [9, 10]. Cells in these and other natural environments experience surface ligand profiles with varying concentration peaks, non-continuity, and large dynamic range [8, 15], differing strongly from smoothly-varying, purely-diffusive environments.

Modern signal processing theory shows that sensing strategies must adapt to the statistics of the input signals, suggesting that spatial sensing in cells should be adapted to the spatial structure of signaling molecules in the cells’ native environments [16]. For example, when designing electronic sensor networks sensing spatial phenomena, adapting sensor placement to the spatial statistic of the signal can significantly improve information acquisition [17]. Furthermore, spatial navigation where sensing plays a key role may also benefit from sensor placement adaptation, as suggested by work from both robot and insect navigation [18, 19]. For example, when navigating turbulent plumes, locusts actively move their antennae to odorant locations to acquire more information on source location [19]. In the context of cell navigation, interstitial gradients can potentially trap cells in local concentration peaks [3]. Cells that can adapt sensing to the patchy structure of the gradient may overcome local traps.

Traditional approaches to studying cell sensing often use highly simplified environmental models, where signals are either uniform or monotonic, neglecting the complex spatial structure in natural cell environments [20–23]. Classic work, beginning with the seminal paper by Berg and Purcell (1977), studied cell sensing within homogeneous environments [20]. This and subsequent works were extended to study the detection of spatially-varying concentrations, where monotonic gradients remain the canonical environmental model [21–23]. Recent work has started to address spatial complexity [24], but much work remains to understand how cell sensing strategies are affected by natural signal distributions, particularly spatially-correlated fluctuations. Such complexity can pose challenges to cell engineering applications, such as CAR-T cell responses to tumor microenvironments [25]. Fundamentally, it is not clear what sense and response strategies are well-adapted to operate in environments where signals take on complex spatial structures.

Interestingly, empirical observations suggest that cells might modulate the placement of their surface receptors to exploit the spatial structure of ligand distribution in its environment [26–33]. For example, some axon guidance receptors, such as Robo1 and PlxnA1, can dynamically rearrange on the surface of growth cones [26, 27]. In such cases, receptors constantly rearrange, adjusting local surface densities in response to changes in ligand distribution across the cell surface. Some chemokine receptors in lymphocytes, such as CXCR4 and CCR2, exhibit similar spatial dynamics [28–30]. Disrupting dynamic rearrangement of CCR2 on the surface of mesenchymal stem cells, without changing its expression level, severely inhibits targeted cell migration to damaged muscle tissues [33]. However, other chemotactic receptors (such as C5aR on the surface of neutrophils) remain uniform even when their ligands are distributed non-uniformly [34]. In addition, during antigen recognition, T-cell receptors (TCRs) take on different placements, ranging from uniform to highly polarized, depending on the density of antigen molecules on the surface of the opposing cell [35]. Thus, across a diverse range of cell surface receptors, we see different, even contradictory rearrangement behavior in response to changes in environmental structure. It remains unclear whether dynamic receptor rearrangement has an overarching biological function across disparate biological contexts.

Inspired by previous works that applied information maximization principles to understand the design of biological systems for signal processing [36–43], we formulate an information-theoretic framework and show that spatial localization of cell surface receptors is an effective spatial sensing strategy in natural cell environments, but relatively inconsequential in purely diffusive environments. Our framework allows us to solve for receptor placements that maximize information acquisition in natural environments, while generating such environments using existing computational models of tissue and soil microenvironments. We find that anisotropic receptor dynamics previously observed in cells are nearly optimal. Specifically, information acquisition is maximized when receptors form localized patches at regions of maximal ligand concentration. Optimizing receptor placement offers a significant gain in information acquisition over uniformly distributed receptors, but only in natural cell habitats, leading to an average of ∼ 1 bit of information gain in tissues and soils but only ∼ 0.01 bits in purely diffusive gradients. The optimal strategy maximizes information by taking advantage of patchy ligand distribution in natural environments, reallocating sensing resources to a small but high signal region on the cell surface, while explicitly “ignoring” ligand information at low signal regions.

Our framework extends naturally to produce a dynamic protocol for continuously redistributing receptors across the cell surface in response to a dynamic environment. We show through simulation that a simple feedback circuit implements this protocol within a cell, redistributing receptors in a signal-dependent manner, and in doing so significantly improving cell navigation. Compared to cells with uniform receptor placement, cells with this circuit achieve more than 30-fold improvement in their ability to localize to the peak of simulated interstitial gradients. Furthermore, our model accurately predicts spatial distributions of membrane receptors observed experimentally [26–30]. Importantly, our framework easily extends to study how spatial organization of many different cellular components, beyond receptor placement, affects information processing (see Discussion). Taken together, our model serves as a useful conceptual framework for understanding the role of spatial organization of signal transduction pathways in cell sensing, and provides a sensing strategy that is both effective in natural cell environments and amenable to cell engineering.

## Results

### An optimal coding framework allows the computation of optimal receptor placement given spatial signal statistics

We are interested in optimal strategies for a task we refer to as spatial sensing. Spatial sensing is an inference task where a cell infers external profiles of varying ligand level across its surface from an internal profile of varying receptor activity across its membrane. This task is a useful model since optimizing performance on this task should improve the cell’ s ability to infer diverse environmental features.

We developed a theoretical framework to study whether manipulating the placement of cell surface receptors can improve spatial sensing performance. Optimizing spatial sensing by tuning receptor placement is analogous to optimizing distributed electronic sensor network by adjusting the location of sensors, which has been extensively studied in signal processing [17]. In the optimization of distributed sensor networks monitoring spatial phenomena (Figure 1A), it is well-known that adjusting the placement of a limited number of sensors can significantly boost sensing performance, where the optimal placement strategy is dictated by the statistics of the input signals [17, 44]. The collection of a limited number of receptors on the cell surface also functions as a distributed sensor network, sensing a spatial profile of varying ligand concentration (Figure 1A). Therefore, we hypothesized that receptor placement can be tuned to improve spatial sensing, and that the optimal strategy depends on the statistics of ligand profiles that cells typically encounter. Unlike traditional works in sensor optimization which focuses on finding a single “best” placement [17], cells can rearrange their receptors within a matter of minutes [27], leading to a potentially much richer class of strategies.

**Figure 1:**
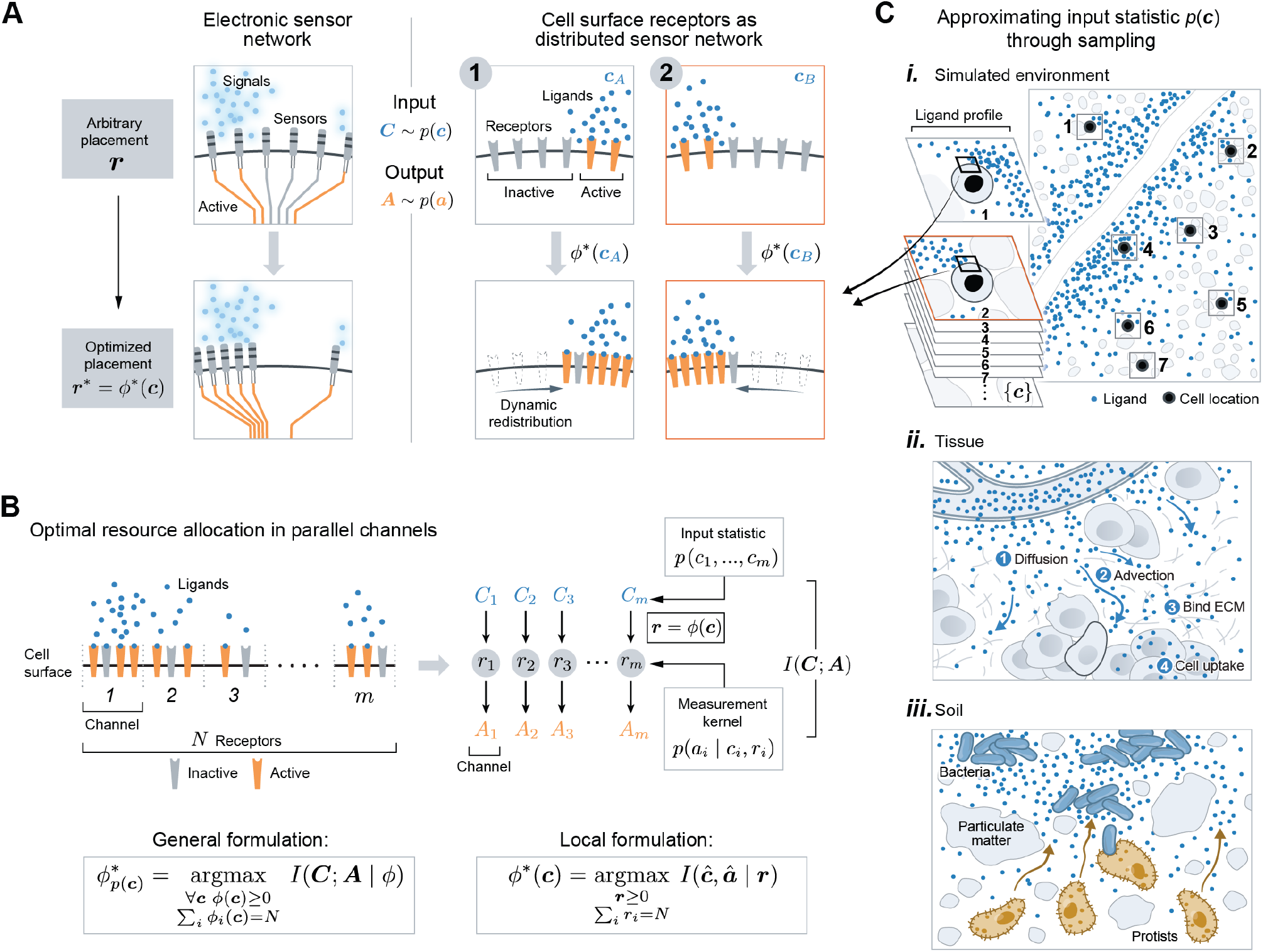
Adapting receptor placements to signal (input) statistic of natural cell environments. (A) (Left) tuning sensor placement can boost the performance of electronic sensor network. (Right) cell surface receptors also function as a sensor network, taking as inputs ligand profiles ***C*** across the cell surface and producing as outputs a profile of receptor activity ***A*** across the cell membrane. The optimal receptor placement strategy *ϕ*^*^ : ***c*** → ***r*** maps each ligand profile to a receptor placement, such that the mutual information *I*(***C***; ***A***) is maximized. (B) The problem of optimal receptor placement formulated as a resource allocation problem over parallel, noisy communication channels. The *i*-th channel represents the *i*-th region of the cell membrane, with input *C*_*i*_, output *A*_*i*_ and receptor number *r*_*i*_. The input statistic *p*(***c***) depends on the environment, and the measurement kernel *p*(*a*_*i*_ |*c*_*i*_, *r*_*i*_) is modeled as a Poisson counting process. The general formulation of the optimal strategy *ϕ*^*^ allocates *N* receptors to *m* channels for each ligand profile ***c***, such that *I*(***C, A***) is maximized (Equation 2). The local formulation selected the receptor placement *ϕ*^*^(***c***) that maximizes *I*(***ĉ, â***), where ***ĉ*** is a Poisson random vector with mean equal to ***c*** (Equation 4). (C) i. Approximating input statistic by simulating natural environments and sampling ligand profiles {***c***} by tiling cells uniformly across the environment; ii. modeling ligand distribution in tissue microenvironment by incorporating diffusion, advection, ECM binding, degradation, and cell uptakes. iii. modeling ligand distribution in soil microenvironment by generating bacteria distributed in spatial patches, releasing diffusive ligands.

Before presenting the general optimization problem, we set up the mathematical framework through the lens of information theory. Consider a two-dimensional (2D) cell with a 1D membrane surface. By discretizing the membrane into *m* equally-sized regions, we modeled the membrane-receptor system as *m* parallel communication channels (Figure 1B). The input to these *m* channels is ***C*** = (*C*_1_, *…, C*_*m*_), where *C*_*i*_ is a random variable representing the amount of ligands at the *i*-th membrane region. The receptor profile ***r*** = (*r*_1_,.., *r*_*m*_) denotes the amount of receptors allocated to each membrane region. The output ***A*** = (*A*_1_, *…, A*_*m*_) is the amount of active receptors across the membrane, which depends on ***c*** and ***r*** through *p*(***A*** = ***a***|***c, r***), the measurement kernel. Consider a placement strategy *ϕ* : ***c*** → ***r***, mapping a ligand profile to a receptor placement (Figure 1B). For a fixed number of receptors *N*, we are interested in the choice of *ϕ* that maximizes the mutual information *I*(***C***; ***A***) between the channels’ inputs ***C*** and outputs ***A***, defined as,

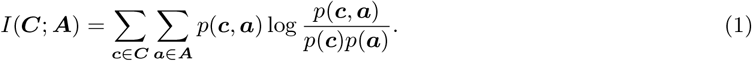

The mutual information quantifies the “amount of information” obtained about ***C*** by observing ***A***. It is minimized when ***C*** and ***A*** are independent, and maximized when one is a deterministic function of the other. All logs are taken in base 2, so information is report in units of bits. Importantly, note the choice of m (membrane bins) sets an upper bound on the mutual information, and hence sets the scale for all information value reported in this paper (see supplement subsection 1.3 for derivations of this relation). Mathematically, the optimal strategy *ϕ*^*^ can be written as

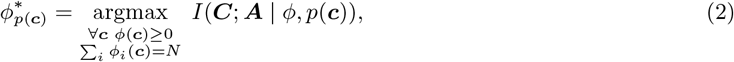

where *N* is the total number of receptors which is taken to be a constant. Note the mutual information converges towards its upper bound as *N* increases (Figure S1A). The mutual information is agnostic to the decoding process in that it does not assume any details about downstream signaling, nor the exact environmental features a cell may try to decode, expanding the scope of our results.

To solve for *ϕ*^*^, we needed to specify both a measurement kernel *p*(***a***|***c, r***) and an input statistic *p*(***c***). We modeled *p*(***a***|***c, r***) assuming that each receptor binds ligands locally and activates independently. Furthermore, each local sensing process is modeled as a Poisson counting process. These assumptions yield the following measurement kernel,

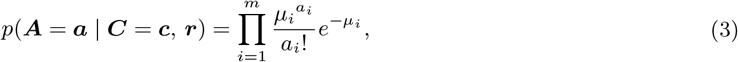

where 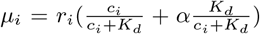 is the average number of active receptors at the *i*-th membrane region. *K*_*d*_ is the equilibrium dissociation constant and *α* accounts for constitutive activity of receptors observed in cells, including many GPCRs, which we take to be small (*α* ≪1) [45, 46]. The bracket term represents the probability of receptor activation, and the fractional term 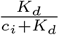 ensures it is always less than 1 [47].

Next, we specify the input statistic *p*(***c***) for three classes of environments: soil, tissue, and monotonic gradient. For each class of environment, we constructed *p*(***c***) empirically, by computationally generating a ligand concentration field as the steady-state solution of a partial-differential equation (PDE), and sampling ligand profiles ({***c***}) from them by evaluating the PDE solution around cells placed at different spatial locations (Figure 1C-i) (for details see Supplement, section 4). Putting the empirical measure on the samples {***c***} approximates the true distribution of ***C***. For soil, we follow mathematical models from [48] and [9], modeling diffusive ligands released from a group of soil bacteria whose spatial distribution agrees with the statistical properties of real soil colonies (Figure 1C-iii, Figure 2A). For tissue, we adopted models from [5] and [49], where they modeled diffusive ligands released from a localized source, perturbed by *in vivo* processes such as interstitial fluid flow and heterogeneous ECM binding, leading to an immobilized interstitial gradient (Figure 1C-ii, Figure 2B). We also considered a monotonic gradient (Figure 2B) as an exponential fit to the simulated interstitial gradient. Fitting ensures any difference between the two environments are due to differences in local structures, not global features such as gradient decay length or average concentration. It is important to note that the overall framework can accommodate any choice of *p*(***c***) and *p*(***a***|***c***) beyond what we have considered.

**Figure 2:**
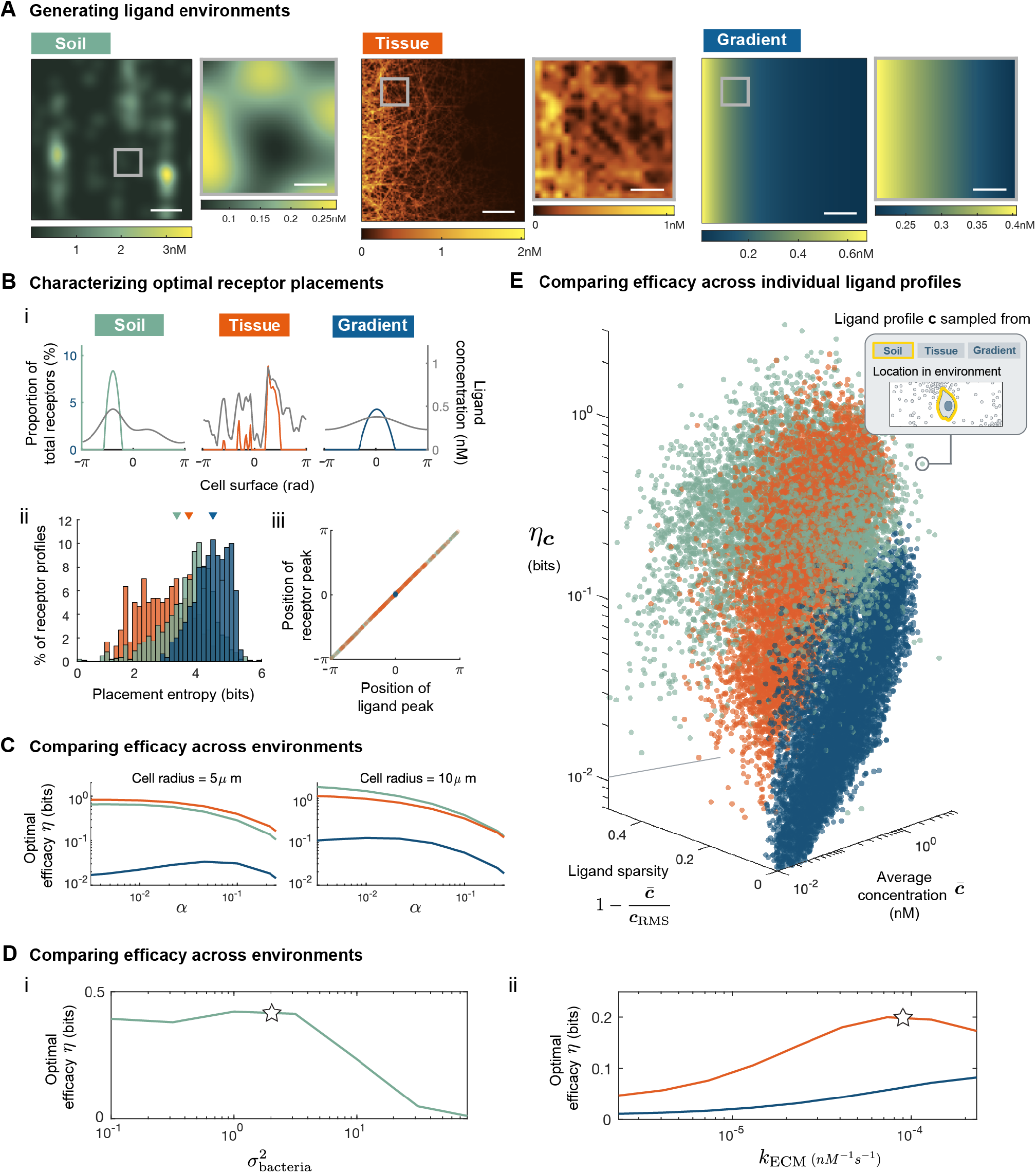
Receptor localization optimizes information acquisition in natural environments. (A), computationally generated ligand concentration fields using PDE models of soil (left), tissue (interstitium) (middle), and simple exponential gradient (right, fitted to tissue with correlation index *R*^2^ = 0.98), all scalebar = 100*μm*, see Table S1 for environment simulation parameters. (B), i) Example of optimal receptor profile *ϕ*^*^(***c***) (colored) and the corresponding ligand profile ***c*** (gray); ii) entropy for each optimal receptor placements in {*ϕ*^*^(***c***)} colored by environment, colored triangles indicate the entropy of three receptor placements shown in i); iii) scatter plot where each dot corresponds to an optimal placement *ϕ*^*^(***c***), x-axis is membrane position with the most receptor, y-axis is membrane position with most ligand in ***c***. (C), optimal efficacy *η* colored by environments, computed with ligand profiles {***c***} sampled using cells of different radius, for receptors of different constitutive activity *α*; see Figure S1B for result with different bin number *m*. (D), i) efficacy for soils environment simulated using different values of 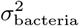, ii) efficacy for tissue environment simulated using different values of *k*_ECM_, and for exponential gradients fitted to each tissue (gradient). Stars correspond to parameter values used to generate panel A-C and E. (E), scatterplot where each dot corresponds to a single pair of ***c*** and *ϕ*^*^(***c***), where ***c*** is sampled from environments as illustrated in Figure 1C-i; *η*_***c***_ is defined in Equation 10. Across all panels, *N* = 1000, *K*_*d*_ = 40*nM, α* = 0.1, *m* = 100 (see Figure S4 and Figure 7B for *η* with other parameters).

We are interested in the functional relationship between ligand profiles {***c***} and their optimal receptor placements {*ϕ*^*^(***c***)}. To this end, we computed the optimal receptor placement for each sampled profile ***c*** individually, reducing the general formulation to a local formulation. Given ligand profile ***c***, random vector ***ĉ*** represents local fluctuations of ***c*** due to stochasticity of reaction-diffusion events. In the case of unimolecular reaction-diffusion processes, it can be shown that ***ĉ*** is a Poisson vector with mean equal to ***c***, solution of the PDE. Therefore, we can solve for *ϕ*^*^(***c***) locally by maximizing the mutual information between ***ĉ*** and the resulting output ***â***,

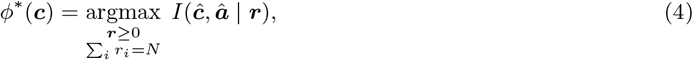

where *p*(***â***) =Σ _***c***_ *p*(***â***_|_ ***ĉ*** = ***c***)*p*(***ĉ*** = ***c***). The main difference between the general formulation of (2) and local formulation of (4) is their dependence on the input statistic *p*(***c***). In the general formulation, the strategy 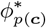 is explicitly parametrized by *p*(***c***). In the local formulation, *ϕ*^*^ is independent of the choice of *p*(***c***). However, differences in *p*(***c***) between environments will still crucially affect the set of optimal receptor profiles that cells will actually adopt, because changing *p*(***c***) changes the region of the domain of *ϕ*^*^ that is most relevant, thus changing the optimal receptor profiles that are actually used in different environments. For example, suppose environment A and B have input statistic *p*_*A*_ and *p*_*B*_, and any ligand profile observed in A is not observed in B, and vice versa. Although *ϕ*^*^ is the same between A and B, this function is being evaluated on entirely different ligand profiles in A compared to B, so that receptor profiles observed in the two environment will likely be very different, in ways dictated by differences between their input statistic *p*_*A*_ and *p*_*B*_. As a result, the statistical structure over the space of ligand profiles plays an important role in determining which receptor placement is effective, even when the placements are computed locally for each ligand profile.

### Receptor localization yields optimal spatial sensing in natural environments

Optimal strategies of receptor placement are similar for soil and tissue environment, where receptors are highly localized within membrane positions experiencing high ligand concentrations. Figure 2B-i shows three examples of optimal receptor placements *ϕ*^*^(***c***) (colored) with the corresponding ligand profile ***c***, one from each class of environments shown in Figure 2A. In all three cases, the peak of each optimal receptor profile is oriented towards the position of highest ligand concentration. Compared to monotonic gradient, receptor profiles optimized for the ligand profiles sampled from tissue and soil are highly localized, with around 80% of receptors found within 10% of the membrane. In general, the optimal strategy consistently allocates more receptors to regions of higher ligand concentration, but in a highly nonlinear manner. Figure 2B-iii shows, across all sampled ligand profiles {***c***}, the peak of receptor profiles always align with the peak of ligand profiles. But instead of allocating receptors proportional to ligand level, receptors tend to be highly localized to a few membrane positions with the highest ligand concentrations.

Indeed, Figure 2B-ii shows that optimal receptor profiles tend to have low entropy. The entropy of receptor profile ***r***, defined as 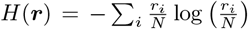, can be used as a measure of localization. Note the maximal value of this entropy measure is limited by the number of membrane bins *m*. Low entropy corresponds to receptor profiles where most receptors are concentrated to a few membrane positions, forming localized patches. Such high degree of localization is partly explained by low receptor numbers. When receptors are limited, information gain per receptor within each membrane channel is approximately independent of receptor number (for details see Supplement, section 2). Thus, the optimal solution allocates all receptors to the channel with the highest information content (see Figure S2). In addition, receptors are more localized for sensing in soil and tissue because locally, they exhibit greater spatial variations in ligand concentration compared to simple gradients (Figure 2A) (for details see Supplement, section 3). Absolute ligand concentration also influences the optimal strategy, which we take to be dilute in agreement with empirical measurements [50, 51]. In saturating environments, the optimal solution completely switches, allocating most receptors to regions of lowest ligand concentrations (for details see Supplement, subsection 3.2, Figure S3). In summary, the optimal placement strategy *ϕ*^*^ in the environments studied can be approximated by a simple scheme, where receptors localize to form patches at positions of high ligand concentration.

Optimally placed receptors significantly improve information acquisition relative to uniform receptors, especially in soil and tissue environments. To make this statement precise, we quantified the efficacy of a receptor placement strategy *ϕ* : ***c*** → ***r*** with respect to a set of ligand profiles {***c***}. First, we denote by *I*_*ϕ*_ the average amount of information acquired by cells adopting the placement strategy *ϕ*,

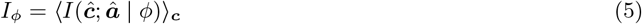

where ⟨·⟩_***c***_ denotes averaging across the set of sample ligand profiles {***c***}, and recall ***ĉ*** is a Poisson-distributed random vector with mean ***c***. In particular, we are interested in information acquisition using the uniform strategy *ϕ*^*u*^ (uniformly distributed receptors) and the optimal strategy *ϕ*^*^,

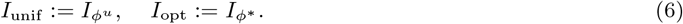

The efficacy of a placement strategy *ϕ* is the change in average information cells acquire by adopting the strategy *ϕ* compared to the uniform strategy,

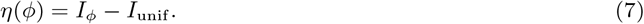

In particular, we are interested in the optimal efficacy,

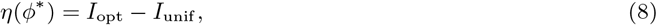

which we will refer to simply as *η* when the dependency is clear from context. For a particular *η*(*ϕ*^*^), the set of ligand profiles {***c***} referred to in its definition is always the same set that *ϕ*^*^ is optimized for. The larger *η* is, the more beneficial it is for cells to place receptors optimally rather than uniformly. We found that *η* is an order-of-magnitude larger for soil and tissue environment compared to a simple gradient (Figure 2C). This difference persists across cells of different size and across a wide range of receptor parameter values (Figure 2C, Figure 7B, Figure S4). In other words, placing receptors optimally rather than uniformly benefits cells in complex, natural environments significantly more than cells in simple, monotonic gradients. Note that differences between tissue and monotonic gradient are due to differences in local spatial structure, not global features such as gradient decay length or global average concentration, as both parameters were made to be identical between the two environments. Lastly, although the mutual information is an exponential measure, so an improvement by one bit has different meaning depending on the baseline *I*_unif_, this fact does not hinder interpretation of *η* as *I*_unif_ is similar between the three environments considered (Figure S4A).

In addition to the large difference in optimal efficacy (*η*) between natural environments and simple gradients, Figure S4 shows similar differences exist when comparing other metrics assessing the benefit of optimizing receptor placement, such as the relative information gain ((*I*_opt_ − *I*_unif_)*/I*_unif_) and the absolute increase in the number of distinguishable input signals 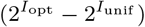. For example, optimizing receptor placement increases the number of distinguishable input states by 40 in tissue, while optimizing the same receptors in the (fitted) gradient environment leads to an increase of 1 state (Figure S4A). Note that in the limit of strong constitutive receptor activity, all placement strategies become equivalent to uniformly distributed receptors. Since receptor activation in the absence of ligands reduces statistical dependence between ligand level and receptor activity, the average information acquisition *I*_*ϕ*_ for any strategy *ϕ* converges to zero for large *α*, driving information gain compared to the uniform strategy to zero (Figure S4A).

For both soil and tissue environment, the optimal efficacy *η* depends on a key parameter in their respective PDE model. We illustrate this dependence by adjusting the value of each respective parameter, sampling new ligand profiles {***c***}, solving for optimal placements {*ϕ*^*^(***c***)}, and computing *η*. Figure 2D shows how *η* changes as we adjust environmental parameters. In soil, *η* drops substantially as ligand sources (bacteria) become more aggregated (Figure 2D-i), corresponding to an increase in the parameter 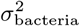 of the random process (see supplement, section 4) used to model bacterial distribution (star corresponds to empirical value from [9]). The decrease in *η* is intuitive since increasing the extent of aggregation of sources makes the environment appear more like a simple gradient generated from a single ligand source. In tissue, optimal efficacy dropped when most ligands were found in solution, instead of bound to the ECM (Figure 2D-ii), corresponding to low ECM binding rate (*k*_ECM_). For reference, star indicates the empirical value of *k*_ECM_ for the chemokine CXCL13 [2]. Compared to its fitted monotonic gradient, *η* in the interstitial gradient remain significantly higher for all ECM binding rates (Figure 2D-ii). In tissue, gradients made up of ECM-bound ligands are ubiquitous, suggesting the optimization of receptor placement is highly relevant.

Optimal efficacy (*η*) is larger in soil and tissue because ligand profiles that cells encounter in such environments tend to be more patchy, having most of the ligands concentrated in a small subset of membrane regions. We make this statement precise by quantifying patchiness of a ligand profile ***c*** using a measure of sparsity,

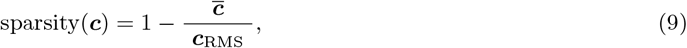

where the root-mean-square 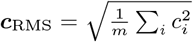 and 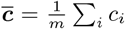 is the average concentration of ***c*** across the membrane. A ligand profile with a sparsity of one has all ligands contained in a single membrane region, whereas a uniform distribution of ligands has a sparsity of zero. Next, we defined an efficacy measure *η*_***c***_ for each ligand profile ***c***,

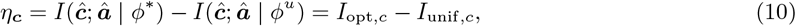

where again *ϕ*^*u*^ denotes uniform receptor distribution. Unlike *η* as defined in Equation 8, *η*_***c***_ does not involve the averaging across the entire set {***c***} through ⟨ · ⟩_***c***_, it measures improvement in information gain for only a single ligand profile ***c***. The larger *η*_***c***_ is, the more useful the optimal placement is for sensing ***c*** compared to a uniform profile. Each dot in Figure 2E corresponds to a ligand profile sampled from an environment, as illustrated in Figure 1C-i. Figure 2E shows that 1) across a wide range of concentrations, sparser ligand profiles tend to induce higher efficacy *η*_***c***_, and 2) ligand profiles sampled from soil and tissue tend to be sparser compared to profiles from the corresponding monotonic gradient. Taken together, since signals cells encounter in natural environments tend to have sparse concentration profiles, cells can improve their spatial sensing performance by localizing receptors to regions of high ligand concentration.

In summary, the value of optimizing receptor placement as a sensing strategy depends strongly on the environmental structure. Patchy ligand distributions found in tissue and soil environments makes optimizing receptor placement a highly effective sensing strategy. Our result demonstrates that uncovering effective cell sensing strategies requires a careful consideration of the spatial structure of the cells’ natural habitat.

### Spatial sensing via the optimal strategy is robust to imprecise placements caused by biological constraints

Despite the optimal strategy *ϕ*^*^ being highly localized and precisely oriented, we found that neither features are necessary to achieve high efficacy. Given the stochastic nature of biochemical processes in cells, this robustness is crucial as it makes the strategy feasible in cells. Fortunately, receptors do not need to adopt *ϕ*^*^ precisely in order to obtain substantial information gain. To illustrate, we perturb the optimal placements and show that sensing efficacy persists when receptors partially align with ligand peak and localize weakly. For soil and tissue, we circularly shift and flatten (by applying a moving average) all optimal receptor profiles {*ϕ*^*^(***c***)} computed from sample ligand profiles to obtain {*ϕ* ^*p*^ (***c***)}, the corresponding set of perturbed profiles. Different degrees of shifting and flattening represents different degrees of misalignment and weakened localization, respectively. Figure 3A shows results of different perturbations (colored) applied to a receptor profile (black). To assess the effect of these perturbations on sensing, we compute the efficacy *η*(*ϕ*^*p*^) of the perturbed profiles, and compare it to the optimal efficacy *η*(*ϕ*^*^). The heatmap in Figure 3B shows the ratio of perturbed to optimal efficacy for various combinations of perturbations, across soil and tissue. Figure 3B-i shows two examples of perturbations (red dots) that drastically alter the receptor profile while still achieving high efficacy. The red and gray curve in the call-out box represents what the “average” perturbed and optimal profiles look like, respectively. They are obtained by circularly shifting each profile in {*ϕ*^*p*^(***c***)} and {*ϕ*^*^(***c***)} so the peak of ***c*** is center, followed by averaging across the set of shifted profiles element-wise. Clearly, highly localized receptors (> 80% of receptors found within 10% of membrane) are not necessary for effective sensing. In fact, compared to uniformly distributed receptors, a modest enrichment of receptors oriented towards the ligand peak (4 folds relative to uniform) already provides significant information gain (Figure 3B). Importantly, enrichment of receptors (around the ligand peak) greater than 4-5 fold has been observed for different membrane receptors [29, 52], and in some cases nearly all receptors redistribute towards the ligand peak [27]. Such robustness holds across different cell sizes and efficacy metric (Figure S5). In tissue, the heatmap of Figure 3B-i also shows that weakly localized receptors (large flatten factor) are more robust to misalignment (large shift factor). Altogether, the robustness results of Figure 3 suggest that biochemical implementations of receptor localization could improve sensing in natural or engineered cells, even in the presence of stochastic fluctuations in a biological circuit that induce imperfect localization. Moreover, the magnitude of receptor enrichment (around the ligand peak) previously observed in cells is sufficient to obtain significant information gain [27, 29, 52].

**Figure 3:**
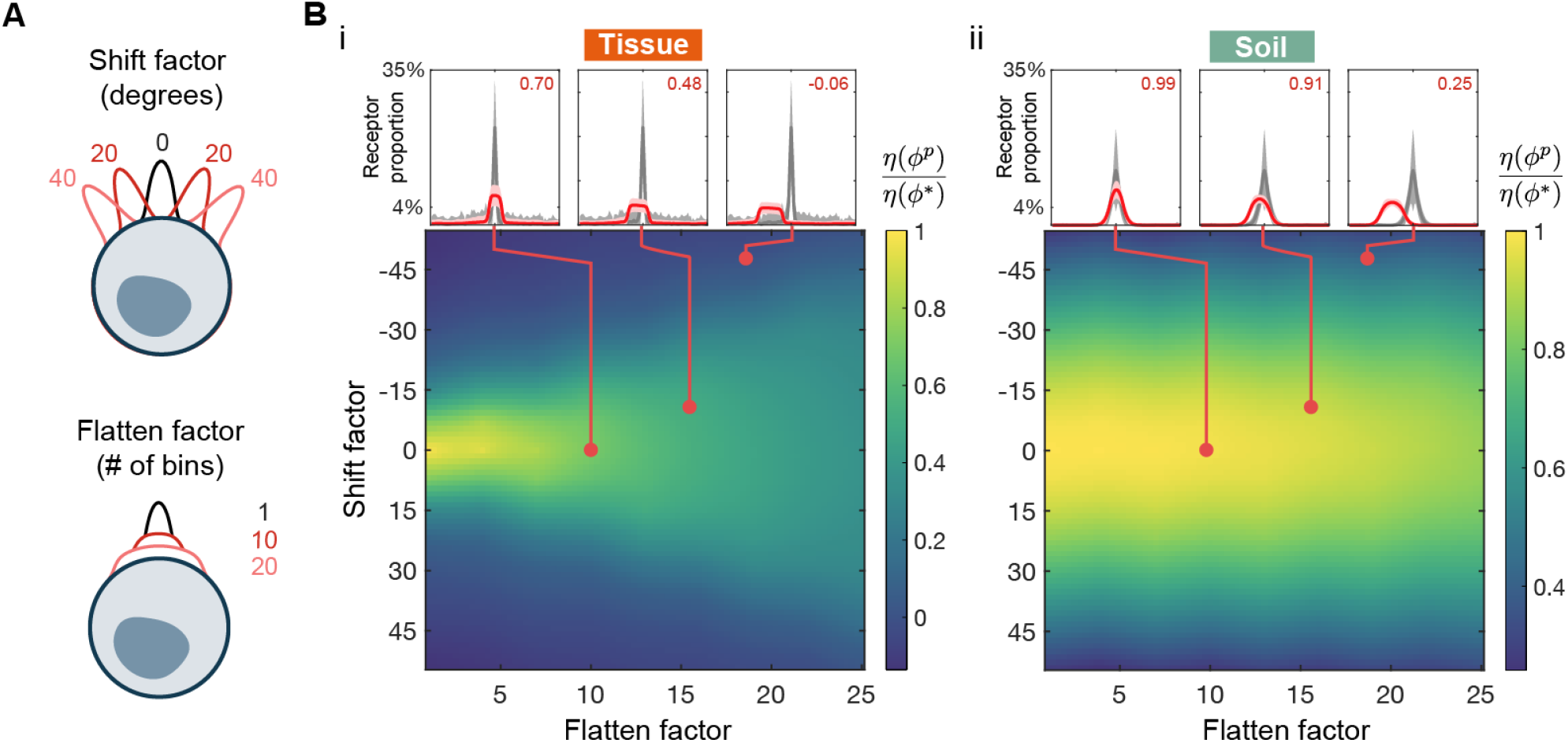
Optimal efficacy *η*(*ϕ*^*^) is robust to minor deviations in receptor placement away from the optimal form. (A), the effect of different degrees of shifting and flattening applied to a receptor profile (black curve). (B), colors of heat map represent ratio of perturbed efficacy *η*(*ϕ*^*p*^) to optimal efficacy *η*(*ϕ*^*^) for different combinations of shifting and flattening, computed for ligand profiles {***c***} sampled from either soil or tissue; call-out boxes corresponds to different sets of perturbations, showing the average of the optimal {*ϕ*^*^(***c***)} (gray) and perturbed {*ϕ*^*p*^(***c***)} (red) receptor placements, after all ligand profile peaks were centered; red numbers indicate the value on heat map; cell radius = 10 μm.

### Optimization framework extends naturally to produce a dynamic protocol for sensing time-varying ligand profiles

Our framework extends naturally to produce a dynamic protocol for rearranging receptors in response to dynamically changing ligand profiles. So far, we have viewed ligand profiles as static snapshots and considered instantaneous protocols for receptor placement. In reality, cells sense while actively exploring their environment, so that the ligand profile it experiences is changing in time, both due to intrinsic changes in the environment state as well as due to the motion of the cell. As the ligand profile ***c***_*t*_ changes over time, we want the receptor profile ***r***_*t*_ to change in an “efficient” manner to improve information acquisition (Figure 4A). Specifically, we obtain a dynamic protocol by extending our framework to account for both information acquisition and a “cost” for changing receptor location. We quantify the cost of moving receptors using the Wasserstein-1 distance *W*_1_(***r***_*A*_, ***r***_*B*_), which is the minimum distance receptors must move across the cell surface to redistribute from profile ***r***_*A*_ to ***r***_*B*_ (for details see Supplement, section 5). For a cell sensing a sequence of ligand profiles 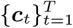 over time, the optimal receptor placement 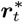 for ***c***_*t*_ now depends additionally on 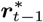, the optimal placement for the previous ligand profile,

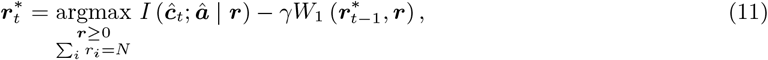

where *p*(***â***) = Σ_***c***_ *p*(***â***|***ĉ***_*t*_)*p*(***ĉ***_*t*_), and *γ* ≥ 0 represents the cost of moving one receptor per unit distance. The cost *γ* implicitly encodes a time scale for receptor redistribution. Smaller *γ* means less “cost” is associated with redistributing receptors, hence the receptor profile becomes more dynamic. The exact relationship between *γ* and the speed of receptor redistribution depends on both receptor properties and the environment, see Figure S6A-D for an example of how receptor speed scales with *γ*. For *γ* = 0, the formulation of Equation 11 reduces to the original static formulation of Equation 4. This dynamic formulation admits a natural interpretation as maximizing information rate (information per receptor-distance moved) instead of absolute information gain. For *t* = 1, we define 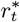 according to the original formulation. Hence, we refer to the dynamic protocol of Equation 11 as the general optimal strategy since it encompasses *ϕ*^*^. Figure 4B illustrates two salient features of this dynamic protocol. Firstly (left), when the peak of the previous receptor profile 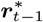 is near the peak of the current ligand profile ***c***_*t*_, 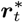 is obtained by shifting receptors towards the current ligand peak but not aligning fully. Secondly (right), when the peak of the previous receptor profile is far from the current ligand peak, some receptors are moved to form an additional patch at the current ligand peak (Figure S6F shows how changing *γ* affects the receptor behavior in Figure 4B). Receptor properties such as the strength of constitutive receptor activity (*α*) also affect receptor redistribution dynamics. For small *α*, receptors localize more readily to align with new ligand peaks because the mutual information term in Equation 11 becomes dominant over the cost of redistribution (Figure S6E). Although the formulation of Equation 11 is quite complex, this general optimal strategy can be achieved by a simple receptor feedback scheme.

**Figure 4:**
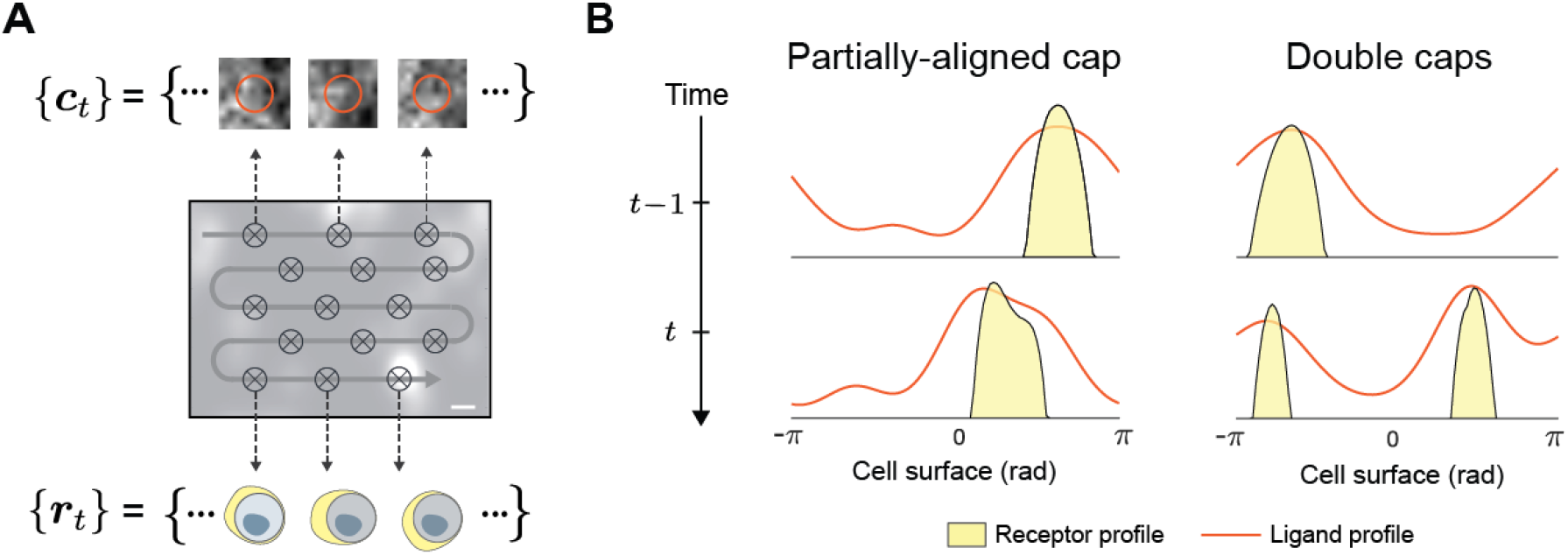
A dynamic receptor placement protocol based on maximizing rate of information gain. (A), schematic showing a cell moving along a path (gray curve) sensing a sequence of ligand profiles {***c***_*t*_} at points (crosses) along the path, using receptor placements 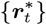 generated by the dynamic protocol. (B), accounting for transport cost, the optimal placement strategy is modified to localize receptors to an intermediate position between subsequent ligand peaks or form multiple receptor peaks.

### Simple feedback scheme rearranges receptors to achieve near-optimal information acquisition

We show that a positive feedback scheme implements the general optimal strategy (Equation 11), dynamically redistributing receptors into localized poles to achieve near-optimal information acquisition. Asymmetric protein localization is a fundamental building block of many complex spatial behaviors in cells, involved in sensing, movement, growth, and division [53]. Many natural localization circuits are well-characterized down to molecular details [54, 55]. Even synthetic networks have been experimentally constructed in yeast, capable of reliably organizing membrane-bound proteins into one or more localized poles [56]. Such works demonstrate the feasibility of engineering new spatial organization systems in cells.

Using a PDE model of receptor dynamics, we show that simple, local interactions can redistribute receptors to achieve near-optimal information acquisition, for both static and dynamic signals. The core feedback architecture of our circuit design uses similar motifs as have been demonstrated in existing synthetic biology circuits [56]. Figure 5A illustrates the three mechanisms (arrows) in our feedback scheme that affects receptor distribution (*r*), which can be expressed mathematically as

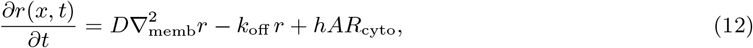

where *x* denotes membrane position and *t* denotes time. The first term represents lateral diffusion of receptors on the membrane with uniform diffusivity *D*. The second term represents endocytosis of receptors with rate *k*_off_. The last term represents incorporation of receptors to membrane position *i* from a homogeneous cytoplasmic pool (*R*_cyto_) with rates *hA*_*i*_, where *h* a proportionality constant and *A*_*i*_ is the local receptor activity (for details see Supplement, section 6), including how parameter values were derived from literature). Activity-dependent receptor recruitment provides the necessary feedback to enable ligand-dependent receptor redistribution. Recent works suggest activity-dependent receptor recruitment can be achieved through biased docking and fusion of secretory vesicles that carry the receptors, to regions of high receptor activity [52, 54, 57]. Budding yeasts Ste2 receptors achieve feedback using an interacting loop with intracellular polarity factor Cdc42 [54]. Note that our feedback scheme is only meant to illustrate one possible implementation of the dynamic rearrangement protocol. Feasible alternatives such as activity-dependent endocytosis or microtubule-dependent receptor redistribution have also been proposed, providing a range of biochemical strategies for implementation of the optimal placement strategy [27, 58].

**Figure 5:**
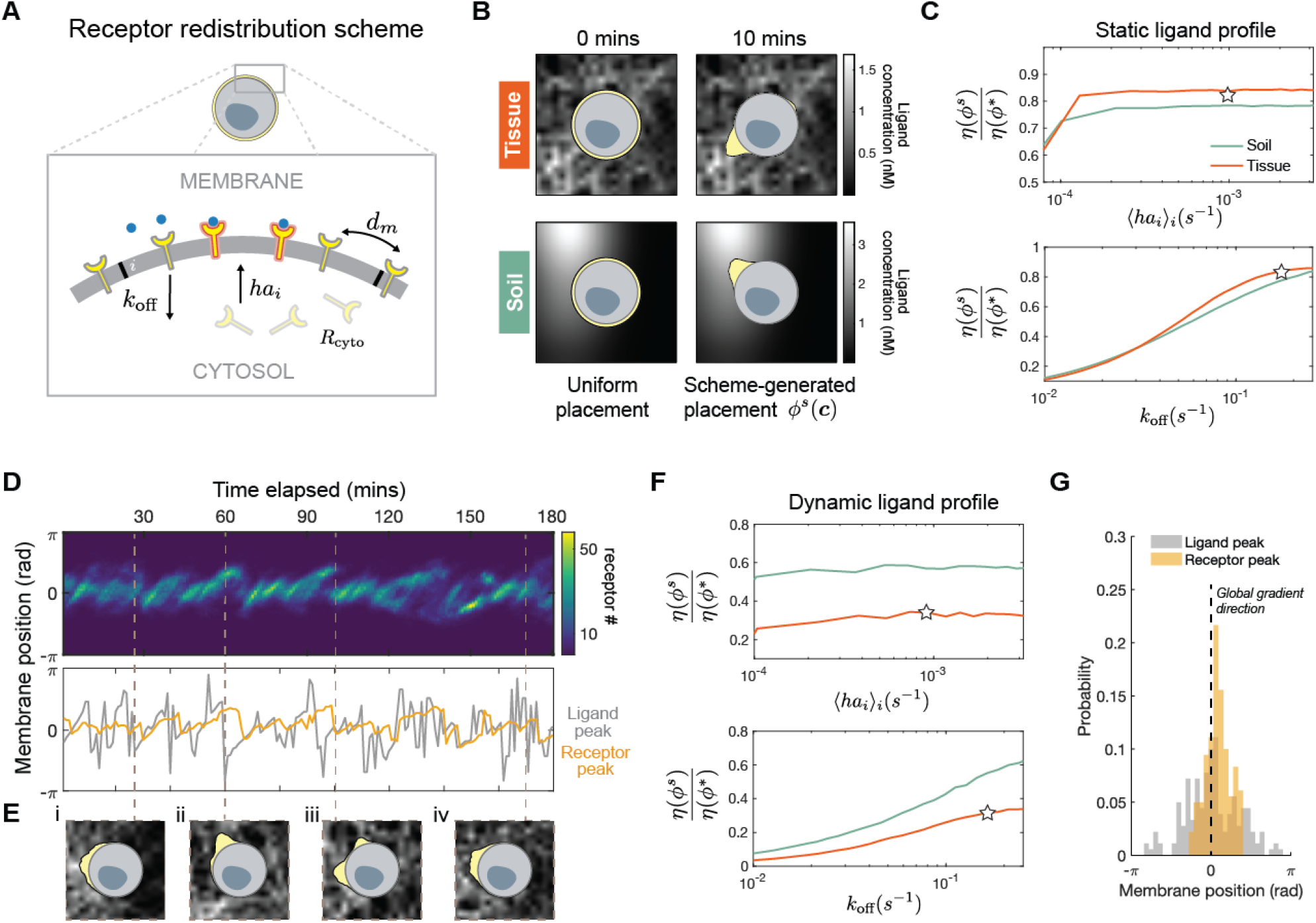
Positive feedback scheme redistributes receptors to achieve near-optimal sensing efficacy for both static and dynamic signals. (A), the cell is modeled as a one-dimensional membrane lattice with a well-mixed cytosol. Receptors are subject to three redistribution mechanisms: endocytosis (*k*_off_), activity-dependent incorporation into membrane (*hA*_*i*_*R*_cyto_), membrane diffusion (*d*_*m*_); the value of *h* sets the feedback strength between receptor activity and the rate with which receptors incorporate into the membrane; *h* = 4 × 10^−3^ s^−1^, *d*_*m*_ = 1 × 10^−2^ μm^2^ s^−1^, *k*_off_ = 1 × 10^−1^ s^−1^ (see supplement, Table S2). (B), receptor profiles (yellow) generated by simulating the feedback scheme for an initially uniform set of receptors, against a static ligand profile from tissue and soil. (C), ratio of scheme efficacy *η*(*ϕ*^*s*^) to optimal efficacy *η*(*ϕ*^*^) for static signals {***c***} sampled from soil and tissue, stars indicate parameter values used for simulation in panel B. (D), (top) kymograph showing the entire temporal sequence of receptor profiles of a moving cell; (bottom) position of ligand peak aligned in time with position of receptor peak as generated by the feedback scheme. (E), snapshots of receptor profiles taken at select time points. (F), ratio of scheme efficacy *η*(*ϕ*^*s*^) to optimal efficacy *η*(*ϕ*^*^) for a sequence of signals {***c***_*t*_} sampled by translating a cell through soil and tissue environment, stars indicate parameter values used for simulation in panel D-E; cell radius = 10*μm* (see Figure S8B-C for results with cell radius = 5*μm*). (G), histogram showing the distribution of ligand peak (gray) and receptor peak (yellow) position on the membrane of the cell from panel D, dashed black line indicates the direction of the global gradient with respect to membrane positions. See Table S2 for feedback scheme simulation parameters.

Given a fixed ligand profile ***c***, Figure 5B shows our feedback scheme can, within minutes, localize receptors (yellow) towards the position of maximum ligand concentration. The rapid localization is robust to changes in *k*_off_ and *h* across at least an order of magnitude (Figure S8A). We denote the steady-state receptor profile generated by our scheme in response to ligand profile ***c*** as *ϕ*^*s*^(***c***). As Figure 5B shows, scheme-generated profiles are far less localized than their optimal counterpart *ϕ*^*^(***c***). Despite this, Figure 5C shows scheme efficacy *η*(*ϕ*^*s*^) are close to that of the optimal value *η*(*ϕ*^*^). Recall *η*(*ϕ*^*^) measures the absolute increase in average information acquired using optimally-placed instead of uniform receptors. Therefore, the scheme efficacy *η*(*ϕ*^*s*^) makes a similar comparison between scheme-driven and uniform receptors. In Figure 5C, we see scheme efficacy is robust to variations in both endocytosis (*k*_off_) and average membrane incorporation rate (⟨*hA*_*i*_ ⟩ _*i*_), with other parameters fixed to empirical values [59]. Stars represent parameters used to simulate profiles in Figure 5B.

Our feedback scheme (Equation 12) can continuously rearrange receptors in response to changes in ligand profile, exhibiting dynamics similar to the optimal dynamic protocol (Equation 11). Figure 5D-E shows a time-varying receptor profile, generated by the feedback scheme in a cell translating across the tissue environment. In this dynamic setting, the scheme can still induce asymmetric redistribution of receptors. Figure 5D (top) shows this dynamic asymmetry through a kymograph of a sequence of receptor profiles {*ϕ*^*s*^(***c***_*t*_)}. As desired, snapshots along this sequence show receptors localize towards regions of high ligand concentration (Figure 5E). Receptor placements generated by our scheme exhibit features of the dynamic protocol shown in Figure 4B. First, as the ligand peak changes position slightly, the receptor peak gets shifted in the same direction after a delay. Figure 5D (bottom) illustrates this phenomena by aligning the time trace of both peak positions. Here, a shift in the ligand peak (gray) is often followed by a corresponding shift in receptor peak (yellow) after an appreciable delay, hence there is only partial peak-to-peak alignment. Second, if the ligand peak changes position abruptly, a second receptor peak forms, oriented towards with the new ligand peak. Figure 5E-iii illustrates this clearly by showing a new receptor peak forming precisely after a large shift in ligand peak position (Figure 5D). We assess the performance of our scheme by comparing scheme-generated placements {*ϕ*^*s*^(***c***)} and optimal placements {*ϕ*^*^(***c***)} corresponding to the same sequence of ligand profiles {***c***_*t*_}. Figure 5F shows that for cells moving in soil and tissue, scheme efficacy *η*(*ϕ* _*s*_) (star) is not far from the optimal value *η*(*ϕ*^*^). Furthermore, scheme efficacy is robust to variations in endocytosis (*k*_off_) and average incorporation rate (⟨*ha*_*i*_ ⟩_*i*_). Taken together, our feedback scheme organizes receptors to achieve near-optimal information acquisition, in both static and dynamic environments.

Our feedback scheme can align receptors with the global gradient direction, suggesting that this scheme may allow cells to escape local ligand concentration peaks within interstitial gradients. On the one hand, Figure 5G shows that the peak of ligand profiles (gray), as experienced by cells, do not always agree with the direction of the global gradient (dashed line) – a known feature of interstitial gradients [3]. On the other hand, receptors organized by the feedback scheme (yellow) align very well with the global gradient direction. The effect of the feedback scheme comes from its ability to localize receptors and account for past receptor profiles. The latter allows the current receptor profile to carry memory of past ligand profiles that the cell has encountered, enabling a form of spatial averaging over ligand peaks. Alignment of receptors to the global gradient should provide significant boost to cell navigation performance, especially in non-monotonic, interstitial gradients.

**Figure 6:**
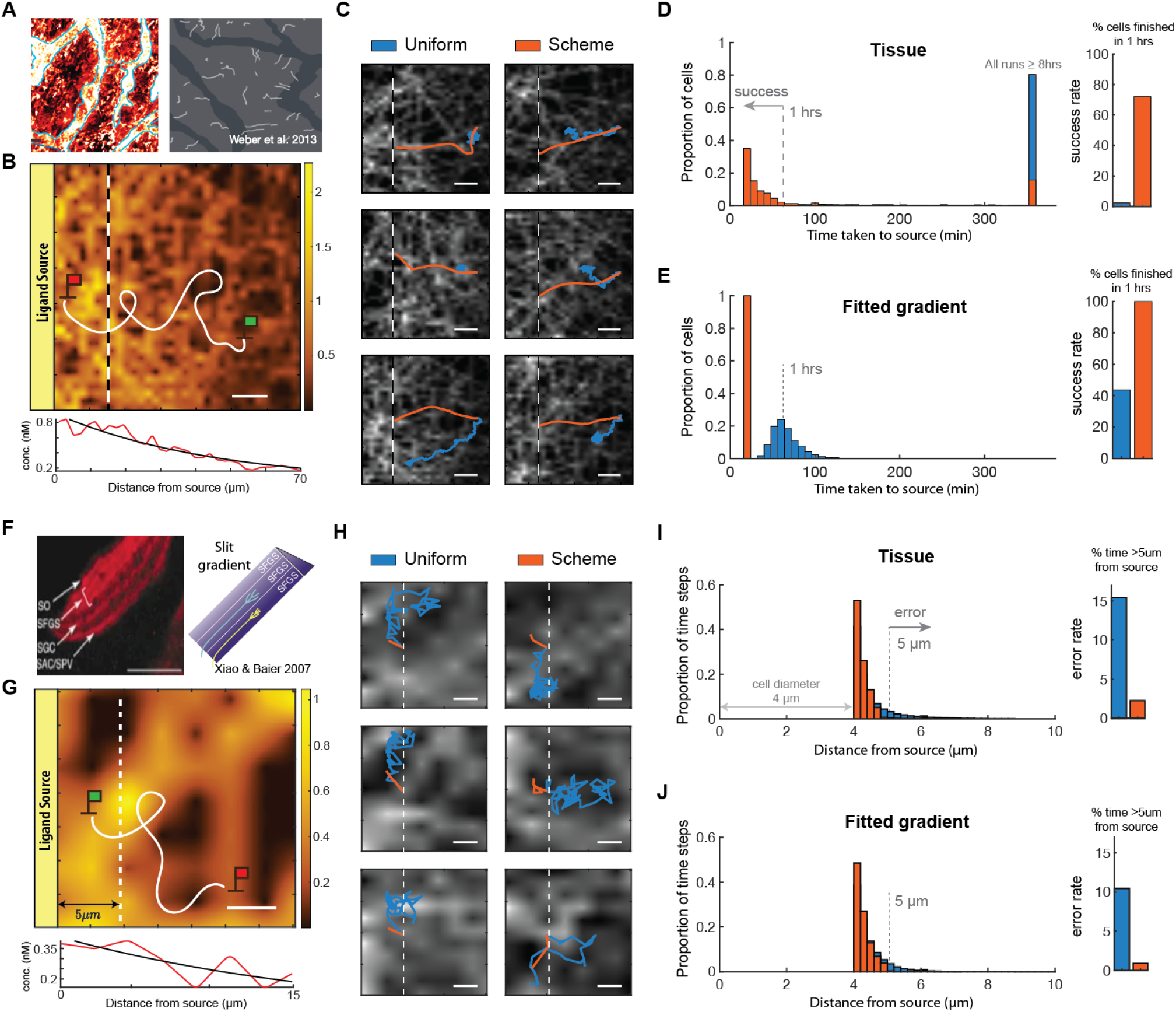
In simulated interstitial gradient, cells localize to source quickly and precisely when receptors are redistributed by the feedback scheme instead of uniformly distributed. (A), (left) interstitial CCL21 gradient, (right) white curves represent haptotactic trajectories of dendritic cells [3]. (B), (top) schematic of a navigation task where a cell (green flag) in a region of an interstitial gradient move towards the source (red flag) by sensing spatially-distributed ligands and decoding source direction locally, the ligand field shown is a region of the tissue environment in Figure 2A obtained through PDE simulation (see supplement section 4); (bottom) red curve shows the tissue ligand field averaged over the y-direction, and black curve is the fitted exponential gradient, scale bar: 10 μm. (C), sample trajectories of repeated simulations of cells navigating with uniform receptors (blue) and with scheme-driven receptors (orange), all scale bars: 10 μm. (D), (left) histogram of time taken to reach source across 600 cells at different starting positions of equal distance from source, note the rightmost bar includes all cells that did not reach the source after 8 hours; (right) barplot showing percentage of runs completed in 1 hrs (success rate), see also Figure S9 for success rate across different simulation parameters. (E), same type of data as in panel D for cells navigating in an exponential gradient (fitted to the interstitial gradient used to generate panel D). (F), red stripes (left) represent growth cones moving within specific lamina along a Slit gradient (right schematic), an ellipse-shaped cell used for this simulation to mimic navigating growth cone, scale bar: 40 μm [60]. (G), (top) schematic of a navigation task where a cell (green flag) senses its environment in order to remain close to source, solid white line represents cell trajectory, dotted white line demarcates a distance of 5 μm from ligand source (see Table S3 for tissue simulation parameters), (bottom) red curve shows the tissue ligand field averaged over the y-direction, and black curve is the fitted exponential gradient, scale bar: 2 μm. (H), sample trajectories of repeated simulations of cells performing task with either uniform or scheme-driven receptors, all scale bars: 2 μm. (I), (left) histogram of time spent by cell at various distance from the ligand source (measured from source to farthest point on cell, perpendicular to source edge) aggregated across 600 cells starting at different positions, moving at 2 hours near the ligand source; (right) barplot shows percentage of time spent more than 5 μm from source (error rate), see also Figure S9 for error rate across different simulation parameters. (J), same data type as panel I for cells navigating in an exponential gradient (fit to interstitial gradient of panel I).

### Feedback scheme enables cells to search quickly and localize precisely in simulated interstitial gradients

Cells using our feedback scheme effectively localizes to the ligand source of simulated interstitial gradients, while cells with uniform receptors become trapped away from the source by local concentration peaks. Immune cells can navigate towards the source of an interstitial gradient in a directed, efficient manner (Figure 6A) [3]. Efficient navigation can be difficult in complex tissue environments, partly due to the existence of local maxima away from the ligand source, potentially trapping cells on their way to the source (Figure 6B). By simulating cell navigation using standard models of directional decoding (for details see Supplement, section 7), we found that cells with uniform receptors can indeed become trapped during navigation. Figure 6C demonstrates this behavior through the trajectories of individual cells with uniform receptors (blue), as they consistently become stuck within specific locations of the environment. On the other hand, using the same method of directional decoding, cells with scheme-driven receptors (orange) reliably reach the source in an efficient manner. Figure 6D illustrates this difference through a histogram of the time it took for a cell to reach the source, created by simulating cells starting at uniformly-sampled locations 40 μm from the source, moving at a constant speed of 2*μm/*min. Remarkably, for the circuit parameter values chosen, only 2% of cells (13/600) with uniform receptors reached the source within 1 hour, compared to 73% of cells (436/600) using the feedback scheme, boosting success rate by more than 30-folds. In fact, Figure 6D shows that > 97% of cells with uniform receptors fail to reach the source even after 6 hours, as expected due to being trapped. Improvement in success rate persists across a wide range of scheme parameters (orders-of-magnitude) and directional decoding schemes (Figure S9). We emphasize that the poor performance of cells with uniform receptors is only partially due to inaccuracy associated with decoding local gradients. Indeed, cells that only follow local gradients have trouble finding the global peak (ligand source) in simulated interstitial gradients, as shown by the fact that cells simulated to move precisely along local gradient directions (direction of maximal increase in ligand concentration across the cell’ s surface) become trapped at local ligand peaks on their way to the source (Figure S9C). As expected, Figure 6E shows that the difference in performance between uniform and scheme-driven receptors is relatively less pronounced in the simple gradient (black curve Figure 6B line plot) – a 2-fold difference in success rate. We discuss the analogy between our feedback scheme and the infotaxis algorithm [61] in the Discussion section.

Our feedback scheme can also help cells remain within a highly precise region along a chemical gradient. During certain developmental programs, cells must restrict their movements within a region along a gradient in order to form stable anatomical structures. Growth cones demonstrate an extraordinary ability in accomplishing this task. Axon projections of retinal ganglion cells can remain within a band of tissue (lamina) of only 3 − 7*μm* wide, at a specific point along a chemical gradient (Figure 6F) [60, 62]. Figure 6G illustrates how we assess our scheme’ s ability to achieve this level of precision. We initiate a cell at a gradient source and track the proportion of time the cell was more than 5 *μm* away from the source. As the cell moves along the gradient, uneven ligand distribution in the environment can lead the cell to move erroneously away from the source. Figure 6H shows that cells with uniform receptors (blue) can indeed make excursions away from the source. But cells with the feedback scheme (orange) reliably stay close to the source for an extended period of time. We quantify this difference by pooling from 600 trajectories of cells starting at different positions along the source, decoding source direction and navigating for 2 hours. Figure 6I shows the number of time steps the cells collectively spent at specific distances from the source. For the circuit parameter values chosen, cells with uniform receptors are found more than 5 μm away from the source 15% of the time (22204*/*144000 steps). On the other hand, cells with the feedback scheme do so only 2% of the time (3287*/*144000 steps), a 7-fold reduction in error rate. Difference in error rate persists for a wide range of scheme parameters and directional decoding schemes (Figure S9D,F). Similar improvement in performance is found for cells navigating in fitted exponential gradients (black curve Figure 6G line plot). Figure 6J shows the error rate is reduced by 10-fold from cells with uniform to scheme-driven receptors (10% vs. 1%). This result is intuitive as the gradients used for this task has extremely short decay length (5 μm) to mimic *in vivo* gradients that growth cones encounter. As a result, the fitted exponential becomes similar to the simulated interstitial gradient. Taken together, our feedback scheme is functionally effective in simulated patchy gradients found in tissue, enabling cells to solve common navigation tasks with significantly improved accuracy and precision.

### Optimal efficacy accurately predicts experimental observations of membrane receptor distribution

In addition to generating optimal sensing strategies for simulated environments, our framework can be used to predict receptor distribution of natural cell surface receptors (Figure 7A), using both the environmental structure in which the receptors function and their biological properties. In addition to environmental structure, receptor properties such as cell surface expression level (*N*) and binding affinity (*K*_*d*_) also play a role in determining the optimal strategy by affecting the measurement kernel (Equation 3). For a simulated tissue environment, Figure 7B shows that despite offering significant information gain (*η* ∼ 1) for a wide range of *N* and *K*_*d*_ values, optimizing receptor placement offers nearly zero gain in information (*η* ≪ 10^−2^) when *N/K*_*d*_ is large. High *N* and low *K*_*d*_ improve information acquisition by allowing the receptor activities to be more sensitive to changes in input level, and since the total amount of information available to the cell is fixed, the amount of additional gain that can be made by optimizing receptor placement is reduced.

**Figure 7:**
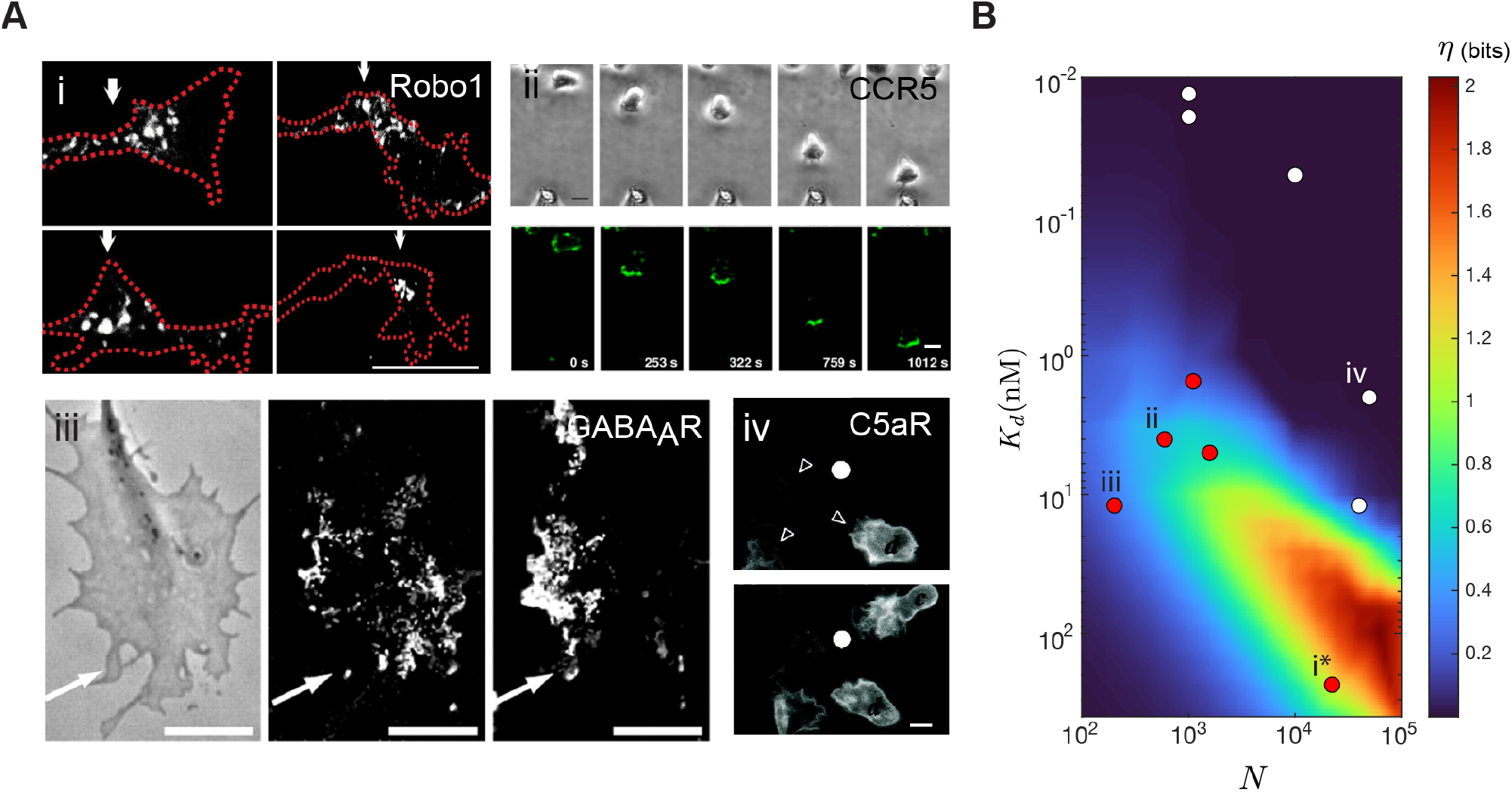
Optimal efficacy *η* predicts observed distributions of cell surface receptors using their surface expression level and binding affinity. (A) observed membrane distributions of receptors in heterogeneous environments, i. white arrowheads indicate Slit receptor Robo1 of commissural growth cones navigating in an interstitial Slit gradient [26]. ii. Human T lymphocytes migrating towards a CCL5-loaded pipette (bottom edge of each panel), top row of panels show brightfield images of a cell taken at different time, bottom row show the corresponding fluorescence images of GFP-tagged CCR5 on the cell surface (time stamp at lower right corner of panel) [63], iii. Effect of a GABA gradient on the distribution of GABA_A_Rs in a growth cone (GC). The arrow indicates direction of the source. (left) Transmission image of GC. (center, right) Images show individual quantum dots-tagged GABA_A_Rs detected by their fluorescence, recorded during the first 10 min of stimulation (center) and during the next 10 min (right) [27], iv. C5aR-GFP remains uniformly distributed in response to a point source of a C5aR agonist, delivered by micropipette (white dot), open arrowheads point to leading edges of cells [64]. Scale bars i,iii: 5 μm, ii,iv: 10 μm. (B) optimal efficacy *η* for different values of *K*_*d*_ and *N* ; values computed using the tissue environment, where the ratio between average ligand concentration and *K*_*d*_ is fixed, *α* = 0.1; red dots correspond to receptors that polarize in heterogeneous environments (CCR2, CXCR4, CCR5, GABA_A_R, Robo1), white dots represent receptors that are constantly uniform (IL-2R, TNFR1, TGF*β*R2, CR3, C5aR), roman numerals correspond to receptors in panel A, see Table S4 for receptor data.

Figure 7B suggests that for real cell surface receptors, we may be able to predict their membrane distribution by specifying both their environment and biological parameters (*N, K*_*d*_). Specifically for receptors functioning in tissue, we predict those with parameters that fall within the high *η* regime (Figure 7B) are more likely to adapt the optimal localized distribution. Although data are limited, empirical observations of real receptors agree with this prediction. Comparing data across cell surface receptors from multiple cell types found in human tissue, Figure 7B show that receptors (red dots) with parameters corresponding to large *η* have been observed to localize in non-uniform environments (Figure 7A-i-iii). Importantly, the localized receptors concentrate at the region of the membrane with the highest ligand concentration, consistent with the theoretically optimal strategy. Such localization is clearly illustrated in panels A-ii and A-iii. Figure 7A-ii shows, within 5 minutes, uniform CCR5 redistributes towards the source of CCL5 placed at the bottom edge. Figure 7A-iii shows GABA receptors localize over time to the membrane region experiencing the highest GABA concentration (arrow indicates source direction). Receptors (white dots) with parameters (*K*_*d*_, *N*) corresponding to small *η*, however, are always uniformly-distributed (Figure 7A-iv), even when the environment is non-uniform. Furthermore, although Figure 7B is based on a fixed *α* (constitutive receptor activity), the striking relationship between receptor organization and optimal efficacy *η* holds for values of *α* spanning at least two orders-of-magnitude (Figure S10B). More detailed comparisons between the experimental receptor distributions and the theoretical optimum is unfortunately not possible, because quantitative descriptions of the ligand profiles experienced by the observed cells are not available. This agreement between theory and observations is not meant to imply that evolution optimizes receptor placement. Indeed, there are key caveats such as variations in receptor expression over time and differences between the environments of different receptors. Our theory does, however, provide a framework for studying natural variations in the spatial organization of receptors, such as differences observed between chemotactic receptors in the same T-cell [28].

## Discussion

A rich collection of works, spanning diverse areas including developmental biology, systems biology, and neuroscience, put forth the idea of optimizing mutual information to predict the design of information processing systems in biology [36–43]. For example, information maximization principles have been applied to derive fundamental limits on the fidelity of information transfer in biochemical networks [43, 65]. Inspired by these works, we formulated an information-theoretic framework that enables us to compute effective cell sensing strategies across different environments. We applied the framework to different signaling microenvironments, including tissues and soils, to discover a receptor localization strategy that significantly improves both cell sensing and navigation. More broadly, our work has a series of conceptual and practical implications. Our theory suggests a functional role for spatial organization in cellular information processing, conceptually showing how spatially-organized intracellular components can be used by cells to more accurately infer the state of its external environment, here through sensing and chemoreceptors. Furthermore, our theory conceptually shows how spatial organization of a cell’ s sensing apparatus can actually reflect spatial structure of its environment. Similar results are found in neuroscience, but it is interesting to see how such an efficient coding perspective can help understand spatial organization within a cell. Lastly, our theory has practical consequences for cell engineering. Currently, most synthetic circuits function without spatial modulation and are studied in well mixed compartments. Our work shows how spatial control over synthetic sense and response architectures can provide new strategies for engineering circuits that function in natural environments.

### Adapting framework to optimize other cell properties with respect to environmental statistic

One can easily adapt our framework to understand how variables other than receptor placement affects spatial sensing. Although this work is about optimizing receptor placement, the key quantity being tuned is the spatial distribution of receptor activity, hence our result is relevant to any variable that 1) affects receptor activity and 2) redistributes across space. To illustrate, consider a generalized model of receptor activation,

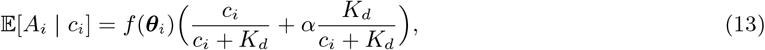

where *f* is an unspecified function of an arbitrary set of variables ***θ***_*i*_, and *f* (***θ***_*i*_) represents the “effective” number of receptors at position *i*. In this work, we considered the case where *θ*_*i*_ = *r*_*i*_ and *f* (*r*_*i*_) = *r*_*i*_, but other factors such as phosphorylation level and membrane curvature also affect local receptor activity *A*_*i*_ [66]. In this way, one can optimize spatial sensing by tuning variables other than receptor placement, by specifying alternative forms of *f*. For example, it is known that given uniformly distributed receptors, those found in membrane regions of higher curvature can exhibit higher activity [66]. Suppose we want to know the optimal way to adjust cell shape to maximize information acquisition, by assuming a linear relationship between local curvature *β*_*i*_ and “effective” receptor number, i.e. *f* (*r*_*i*_, *β*_*i*_) = *β*_*i*_*r*_*i*_. Given uniform receptors and a constraint on total contour length (or area) of the membrane, we quickly arrive at the optimal solution since this problem is now identical to our original formulation. The optimal strategy is to increase membrane curvature at regions of high ligand concentration, by making narrow protrusions (for details see Supplement, section 9).

### Connection between information acquisition and navigation

We showed that a receptor placement strategy aimed at maximizing information rate can boost cell navigation performance. Since information content increases towards the ligand source, receptors are more likely to move towards the side of the membrane closer to the source rather than away, enforcing movement up gradients. Furthermore, the trade-off between information acquisition and receptor redistribution in Equation 11 can be viewed as combining exploitative and exploratory tendencies, where larger redistribution “cost” favors exploitation. This strategy is similar in principle to the infotaxis algorithm [61], where one can view receptors as “navigating agents”, whose movements guide the cell towards the target. Although the idea is quite intuitive, the exact relationship between navigation and information acquisition requires further investigation. On the one hand, the feedback scheme is most effective in the case of limited sampling of inputs (Figure S9B,E), which suggests maximizing information content indeed helps with navigation. On the other hand, moving receptors to maximize information rate is significantly more effective as a navigation strategy compared to only maximizing absolute information (Figure S7).

### Optimizing spatial organization at different stages of information processing

Optimizing information transmission by organizing effectors in space can happen at all stages of signal processing within the cell, but is likely most effective at the receptor level. The most obvious reason is due to the data processing inequality, which states that post-processing cannot increase information. Therefore, only optimization at the level of receptor activation can increase the total amount of information that is available to the cell. The second reason is due to the “hourglass” topology of cell signaling networks, which represent the fact that a large number of signaling inputs converge onto a small number of effectors internal to the cell [67]. For example, G-protein-coupled receptors, one of the largest group of cell surface receptors, drive downstream signaling through the same G-proteins. This feature makes optimizing spatial organization at later stages of information processing very difficult, since information can be easily lost by diffusion of effector molecules activated by different inputs, which ends up “mixing” different spatial signals.

## Data availability

All analysis, simulation and plotting scripts are openly available at: https://github.com/neonine2/receptor-code. All data generated in this work is openly available at: http://dx.doi.org/10.22002/D1.2149.

## Material availability

This paper did not generate new reagents.

## Acknowledgements

We thank Michael Elowitz, Erik Winfree, and David Sivak for scientific discussions and Dominik Schildknecht, Han Kim, Guruprasad Raghavan, Pranav Bhamidipati, Abdullah Farooq, for feedback on the manuscript, Inna-Marie Strazhnik for illustrations, and Angela Anderson for editorial advice. We also would like to thank Eugenio Marco and Katarzyna Rejniak for technical advice with receptor feedback and tissue simulations, respectively. The authors would like to acknowledge the Heritage Medical Research Institute and Packard Foundation for funding and intellectual support.

## Author Contributions

Conceptualization, Z.W. and M.T.; Methodology, Z.W. and M.T.; Manuscript writing, Z.W. and M.T.; Supervision, M.T.; Funding acquisition, M.T.

