## supplemental text for "Localization of signaling receptors maximizes cellular information acquisition in spatially-structured natural environments"

### Supplemental Information

#### 1. Formulation of optimization problem

In this paper, we developed a theoretical framework to study whether manipulating the placement of cell surface receptors can improve the spatial sensing performance. Optimizing spatial sensing by tuning receptor placement is analogous to optimizing distributed electronic sensor network by adjusting the location of sensors, which has been extensively studied in signal processing [1]. Before presenting the general optimization problem, we set up the mathematical framework through the lens of information theory. Consider a two-dimensional (2D) cell with a 1D membrane surface. By discretizing the membrane into $m$ equally-sized regions, we modeled the membrane-receptor system as $m$ parallel communication channels (Figure 1B). The $i$-th channel takes as input $C_i \in \mathbb{N}_0$, a random variable denoting ligand count at the $i$-th region of the membrane surface. Given $r_i \in \mathbb{N}_0$ receptors, this channel produces as output $A_i \in \mathbb{N}_0$, a number of active receptors that is random due to stochastic nature of receptor activation and randomness in $C_i$. Given $m$ channels representing the entire cell membrane, our model comprised four key mathematical objects: ligand profile $\boldsymbol{C} = (C_1, ..., C_m)$, receptor placement $\boldsymbol{r} = (r_1, .., r_m)$, active receptor profile $\boldsymbol{A} = (A_1, ..., A_m)$, and measurement kernel $P(\boldsymbol{A} = \boldsymbol{a} \mid \boldsymbol{C} = \boldsymbol{c}, \boldsymbol{r})$. The input $\boldsymbol{C} \sim p(\boldsymbol{c})$ is now the entire ligand profile across the cell surface. Each realization $\boldsymbol{c}$ of $\boldsymbol{C}$ has probability $p(\boldsymbol{c})$ of being observed. We explain below how $p(\boldsymbol{c})$ can be constructed to represent statistics of ligand profiles cells naturally encounter (see **Input statistic**). The receptor profile $\boldsymbol{r}$ denotes the number of receptor allocated to each membrane region. The output $\boldsymbol{A} \sim p(\boldsymbol{a})$ is the number of active receptors across the membrane, which depends on $\boldsymbol{c}$ and $\boldsymbol{r}$ through $p(\boldsymbol{a}|\boldsymbol{c}, \boldsymbol{r})$, the measurement kernel. We explain below how this kernel can be modeled (see **Measurement kernel**).

Consider a placement strategy $\phi : \boldsymbol{c} \to \boldsymbol{r}$, that maps a ligand profile to a receptor placement. In our general optimization problem (Figure 1B), we are interested in the choice of $\phi$ that maximizes the amount of information the cell can obtain regarding $\boldsymbol{C}$ by observing $\boldsymbol{A}$, for a fixed number of receptors $N$. Formally, we quantify this information using the mutual information,

$$I(\boldsymbol{C}; \boldsymbol{A}) = \sum_{\boldsymbol{c} \in \boldsymbol{C}} \sum_{\boldsymbol{a} \in \boldsymbol{A}} p(\boldsymbol{c}, \boldsymbol{a}) \log \frac{p(\boldsymbol{c}, \boldsymbol{a})}{p(\boldsymbol{c})p(\boldsymbol{a})}. \tag{S1}$$

The mutual information is minimized when $\boldsymbol{C}$ and $\boldsymbol{A}$ are independent, and maximized when one is a deterministic function of the other. Since $p(\boldsymbol{c}, \boldsymbol{a}) = p(\boldsymbol{a}|\boldsymbol{c}, \boldsymbol{r} = \phi(\boldsymbol{c}))p(\boldsymbol{c})$, each summand in the mutual information will be affected by the choice of $\phi$. Taken together, we arrive at our general formulation of the optimal strategy $\phi^*$:

$$\phi^*_{p(\boldsymbol{c})} = \underset{\substack{\forall \boldsymbol{c}\ \phi(\boldsymbol{c}) \geq 0 \\ \sum_i \phi_i(\boldsymbol{c}) = N}}{\operatorname{argmax}} \; I(\boldsymbol{C}; \boldsymbol{A} \mid \phi,\, p(\boldsymbol{c})), \tag{S2}$$

where $N$ is the total number of receptors. The subscript $p(\boldsymbol{c})$ is meant to emphasize the dependence of the optimal strategy on the input statistics.

To solve for $\phi^*_{p(\boldsymbol{c})}$, we needed to specify both a measurement kernel $p(\boldsymbol{a}|\boldsymbol{c}, \boldsymbol{r})$ and an input statistic $p(\boldsymbol{c})$. The input statistic $p(\boldsymbol{c})$ for an environment represents the probability that a cell in that environment will encounter the ligand profile $\boldsymbol{c}$.

##### *1.1. Measurement kernel*

We model $p(\boldsymbol{a}|\boldsymbol{c}, \boldsymbol{r})$ assuming that each receptor binds ligands locally and activates independently of other receptors. These assumptions allow us to factorize $p(\boldsymbol{a}|\boldsymbol{c}, \boldsymbol{r})$ as follows,

$$P(\boldsymbol{A} = \boldsymbol{a} \mid \boldsymbol{C} = \boldsymbol{c},\, \boldsymbol{r}) = \prod_{i=1}^{m} P(A_i = a_i \mid C_i = c_i,\ r_i). \tag{S3}$$

Each local sensing process involves probabilistic ligand-receptor interaction which can be viewed as a Bernoulli process. In this way, the number of active receptors follows a Binomial distribution, which can be approximated with the Poisson distribution when the probability of successful binding event is low. Indeed, experimental measurements have shown that receptor occupancy is well-approximated by the Poisson distribution [2], such that

$$P(A_i = a_i \mid C_i = c_i,\, r_i) = \frac{{\mu_i}^{a_i}}{a_i!} e^{-\mu_i}, \tag{S4}$$

where $\mu_i = r_i\left(\frac{c_i}{c_i+K_d} + \alpha\frac{K_d}{c_i+K_d}\right)$. The bracket term represents the probability of activation for a receptor experiencing $c_i$ ligands. $K_d$ is the equilibrium dissociation constant and $\alpha$ represents constitutive receptor activity, which we take to be small ($\alpha \ll 1$). In other words, the number of active receptors $A_i$ given ligand count $c_i$ is a Poisson random variable with mean $\mu_i$. Equation (S3) and (S4) together specify the measurement kernel.

*1.2. Input statistic*

Next, we specify the input statistic $p(\boldsymbol{c})$ which will be determined by spatial distribution of ligands, thus differ between different classes of environment. Suppose a circular cell samples its environment by binding nearby ligands. The cell will encounter certain spatial profiles of ligands more often than others, and such statistics will likely depend on the type of environment the cell lives in. In this work, we studied three classes of environments: soil, tissue, and monotonic gradient. Closed form models do not exist for ligand profile statistics of natural environments. Therefore, we take an empirical approach, generating instances of each environment as the steady-state solution of a partial-differential equation (PDE) models and directly sample ligand profiles from them (see section 4 for details on all PDE models). For soil, we adopted mathematical models from [3] and [4], modeling diffusive ligands released from a group of soil bacteria whose spatial distribution agrees with the statistical properties of real soil colonies (Figure 1C-iii, Figure 2A). For tissue, we adopted models from [5] and [6], where they modeled diffusive ligands released from a localized source, perturbed by *in vivo* processes such as interstitial fluid flow, non-uniform ECM binding and cell uptake, to represent an interstitial gradient (Figure 1C-ii, Figure 2B). We also considered a simple (monotonic) gradient (Figure 2C) which is an exponential fit to the simulated interstitial gradient (Figure 2B). Fitting ensures any difference between the two environments are due to differences in local structures, not global features such as gradient decay length or average concentration. For each environment, we obtain a ligand concentration field $c(x)$ as the steady-state solution of a PDE. Then, we tile it with a cell of fixed size and evaluate the concentration field along each cell membrane to obtain a set of ligand profiles denoted $\{\boldsymbol{c}\}$ (Figure 1C-i). Putting the empirical measure on the samples $\{\boldsymbol{c}\}$ approximates the true distribution of $\boldsymbol{C}$. It is important to note that although we modeled $p(\boldsymbol{c})$ and $p(\boldsymbol{a}|\boldsymbol{c})$ in these ways, the overall framework can accommodate any alternative choices of model.

For these choices of $p(\boldsymbol{c})$ and $p(\boldsymbol{a}|\boldsymbol{c})$, we aimed to study the functional relationship between ligand profiles $\{\boldsymbol{c}\}$ and their optimal receptor placements $\phi^*(\boldsymbol{c})$. To this end, we optimized receptor profiles for each sampled profile $\boldsymbol{c}$ individually, reducing the general problem to a local formulation. Given ligand profile $\boldsymbol{c}$, the random vector $\hat{\boldsymbol{c}}$ represents local fluctuations of $\boldsymbol{c}$ due to stochasticity of reaction-diffusion events. In the case of unimolecular reaction-diffusion processes, it can be shown that $\hat{\boldsymbol{c}}$ is a Poisson random vector with mean equal to $\boldsymbol{c}$, solution of the PDE. Therefore, we can solve for $\phi^*(\boldsymbol{c})$ locally by maximizing the mutual information between $\hat{\boldsymbol{c}}$ and the resulting output $\hat{\boldsymbol{a}}$,

$$\phi^*(\boldsymbol{c}) = \underset{\substack{\boldsymbol{r}\geq 0 \\ \sum_i r_i = N}}{\operatorname{argmax}}\; I(\hat{\boldsymbol{c}}, \hat{\boldsymbol{a}} \mid \boldsymbol{r}), \tag{S5}$$

where $p(\hat{\boldsymbol{a}}) = \sum_{\boldsymbol{c}} p(\hat{\boldsymbol{a}}|\hat{\boldsymbol{c}} = \boldsymbol{c})p(\hat{\boldsymbol{c}} = \boldsymbol{c})$ and $N$ is the total receptor number. We assume $\boldsymbol{r}$ to be real-valued instead of integer-valued when solving (S5), this is reasonable as long as $N$ is not too small.

The main difference between the general formulation of (S2) and local formulation of (S5) is their dependence on the input statistic $p(\boldsymbol{c})$. In the general formulation, the strategy $\phi^*_{p(\boldsymbol{c})}$ is explicitly parametrized by $p(\boldsymbol{c})$. In the local formulation, $\phi^*$ is independent of the choice of $p(\boldsymbol{c})$. However, differences in $p(\boldsymbol{c})$ between environments will still crucially affect the set of optimal receptor profiles that cells will actually adopt. This is because changing $p(\boldsymbol{c})$ changes the region of the domain of $\phi^*$ that is most relevant, thus changing the optimal receptors profiles that are actually used in different environments. For example, suppose environment A and B have input statistic $p_A$ and $p_B$ with non-overlapping support, meaning that any ligand profile observed in A is not observed in B, and vice versa. Although $\phi^*$ is the same between A and B, this function is being evaluated on entirely different ligand profiles in A compared to B, so that receptor profiles observed in the two environment will likely be very different, in ways dictated by differences between their input statistic $p_A$ and $p_B$. As a result, the statistical structure over the space of ligand profiles plays an important role in determining which receptor placement is effective, even when the placements are computed locally for each ligand profile.

The constrained nonlinear optimization problem of (S5) was evaluated using the fmincon routine of Matlab 2021 [7]. The Sequential Quadratic Programming algorithm was used to ensure accurate solutions that may exist near the boundary of the feasible region. Furthermore, the analytical gradient of the objective function, shown in equation (S24), was supplied to ensure faster convergence.

##### *1.3. Bin number and mutual information*

An important point to emphasize is that the choice of $m$ (number of discrete membrane bins) sets a scale for all information values reported in the paper, because the mutual information ($I(\boldsymbol{C};\boldsymbol{A})$) is bounded by the entropy of its input which scales logarithmically with $m$. We derive such an upper bound on $I(\boldsymbol{C};\boldsymbol{A})$ by first considering the following general property of mutual information. For any pair of discrete random variables $X$ and $Y$, taking values in $\mathcal{X}$ and $\mathcal{Y}$, respectively, their mutual information $I(X;Y)$ can be equivalently expressed as,

$$\begin{aligned} I(X;Y) &= \sum_{x\in\mathcal{X},y\in\mathcal{Y}} p_{(X,Y)}(x,y)\log\frac{p_{(X,Y)}(x,y)}{p_X(x)p_Y(y)} \\ &= \sum_{x\in\mathcal{X},y\in\mathcal{Y}} p_{(X,Y)}(x,y)\log p_{X|Y=y}(x) - \sum_{x\in\mathcal{X},y\in\mathcal{Y}} p_{(X,Y)}(x,y)\log p_X(x) \\ &= -\sum_{y\in\mathcal{Y}} p_Y(y)\mathrm{H}(X\mid Y=y) - \sum_{y\in\mathcal{Y}} p_X(x)\log p_X(x) \\ &= H(X) - H(X\mid Y). \end{aligned} \tag{S6}$$

Since $X$ and $Y$ are discrete random variables, $H(X\mid Y)$ must be non-negative, which implies,

$$I(X;Y) \le H(X). \tag{S7}$$

Furthermore, suppose $X = (X_1,\ldots,X_m)$ is a discrete, multivariate random variables, then we can bound its entropy $H(X)$ using the fact that the joint entropy of a set of variables is less than or equal to the sum of the individual entropies of the variables in the set,

$$\begin{aligned} H(X) &\le \sum_{i=1}^{m} H(X_i) \\ &\le m\max_i H(X_i). \end{aligned} \tag{S8}$$

Since $\boldsymbol{C} = (C_1,\ldots,C_m)$ and $\boldsymbol{A} = (A_1,\ldots,A_m)$ are both discrete (multivariate) random variables, Equation S7 and Equation S8 both apply and we immediately get the following bound on the mutual information $I(\boldsymbol{C};\boldsymbol{A})$,

$$I(\boldsymbol{C};\boldsymbol{A}) \le m\max_i H(C_i), \tag{S9}$$

where $m$ is the number of membrane bins. To simplify Equation S9 further, we will need to consider a specific sensing environment. Let us consider a cell sensing an average of $\bar{c}$ molecules distributed uniformly across space. In this simple environment, the number of ligand molecules at each of the $m$ membrane bin are identically represented by a Poisson random variable with mean $\bar{c}/m$,

$$C_i \sim \text{Pois}(\bar{c}/m), \quad i = 1, \ldots, m. \tag{S10}$$

Since all components $C_i$ are now identically distributed, Equation S9 reduces to,

$$I(\boldsymbol{C}; \boldsymbol{A}) \leq mH(C_i), \tag{S11}$$

where $C_i \sim \text{Pois}(\bar{c}/m)$. The entropy of a Poisson random variable $C_i$ with parameter $\lambda$ takes on the form,

$$H(C_i) = \lambda[1 - \log(\lambda)] + e^{-\lambda} \sum_{k=0}^{\infty} \frac{\lambda^k \log(k!)}{k!}, \tag{S12}$$

where $\lambda = \bar{c}/m$. Combining Equation S11 and Equation S12 gives the bound,

$$\begin{aligned} I(\boldsymbol{C}; \boldsymbol{A}) &\leq mH(C_i) \\ &\leq m \left( \frac{\bar{c}}{m}[1 - \log(\bar{c}/m)] + e^{-\bar{c}/m} \sum_{k=0}^{\infty} \frac{(\bar{c}/m)^k \log(k!)}{k!} \right) \\ &\leq \bar{c}[1 - \log(\bar{c}/m)] + e^{-\bar{c}/m} \sum_{k=2}^{\infty} \frac{\bar{c}^k \log(k!)}{k! m^{k-1}} \\ &\leq \bar{c}[1 - \log(\bar{c}/m)] + \frac{1}{m} \sum_{k=2}^{\infty} \frac{\bar{c}^k \log(k!)}{k!}, \end{aligned} \tag{S13}$$

where the second term goes to zero as $m$ goes to infinity since the infinite sum converges. For large $m$, therefore, we obtain an upper bound on the mutual information $I(\boldsymbol{C}; \boldsymbol{A})$ that scales logarithmically with the number of membrane bins $m$,

$$I(\boldsymbol{C}; \boldsymbol{A}) \leq \bar{c}(1 - \log(\bar{c})) + \bar{c}\log(m). \tag{S14}$$

Figure S1A shows this upper bound (red) for $\bar{c} = 1$. As further validation of Equation S14, Figure S1A shows that as we increase receptor number $N$, $I(\boldsymbol{C}; \boldsymbol{A})$ converges toward the derived upper bound. Furthermore, the result that optimizing receptor placement is significantly more beneficial in natural environments compared to simple gradients holds for a wide range of membrane bin numbers, as shown in Figure S1B where the absolute information gain ($\eta$, Equation 8) is significantly larger in natural environments compared to simple gradients, for a wide range of $m$ values.

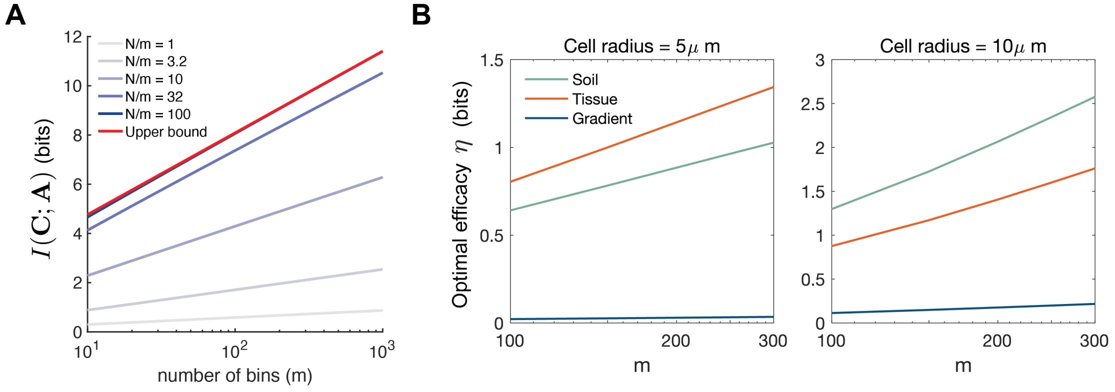


Figure S1: **Effect of the number of membrane bins ($m$) on mutual information and optimal efficacy, Related to Figure 1 and 2** (A) The maximum mutual information achievable (red line, Equation S14) as a function of m, the number of membrane bins. As the number of receptors per bin ($N/m$) increases, the mutual information converges to its maximum value. (B) the optimal efficacy for different choices of $m$, computed across tissue, soil, and simple gradient, $\alpha = 0.01, K_d = 40, N = 1000$.

#### 2. Theoretical properties of Poisson channels

In information theory, the Poisson channel is a canonical model used to study communication of information by random discrete occurrences in time that obey Poisson statistics. We show that we can map our receptor activation model directly onto this canonical model. As a result, we make use of existing results from information theory regarding the Poisson channel to 1) show that the localized receptor placement strategy described in the main text holds across most "reasonable" biochemical models of receptor activation, and 2) provide intuition for how different factors such as ligand concentration can alter the optimal strategy.

##### *2.1. Mapping receptor model to the canonical Poisson channel model*

We begin by showing how a single membrane receptor channel can be mapped to the canonical scalar Poisson model. The same argument applies for mapping multiple parallel membrane receptor channels to the canonical vector Poisson model introduced in the next section.

Recall the receptor model of Equation S4 we used to represent the number of active receptors $A$ for a given ligand level $\boldsymbol{c}$, which is motivated by empirical measurements of receptor activity,

$$p(A = a \mid C = c, r) = \frac{\mu^a}{a!}e^{-\mu}, \;\; \mu = r(\frac{c}{c + K_d} + \alpha\frac{K_d}{c + K_d}). \tag{S15}$$

Although this model of receptor activation consists of many biochemical details, we can map it directly onto the canonical scalar Poisson model,

$$Y|X \sim \text{Pois}(\alpha X) \tag{S16}$$

where $X$ is a scalar input, $Y$ is a scalar output, and $\alpha$ is a scaling variable. Such a model defines a Poisson channel whose output is a Poisson random variable conditioned on the input $X$ with its mean equal to $rX$, where $X$ is an arbitrary input random variable. We map this channel model maps onto our model of receptor activation for a single membrane region, by defining $X$, $\alpha$ in the following way,

$$\begin{aligned} X &:= f(C) = \frac{C}{C + K_d} + \alpha\frac{K_d}{C + K_d}, \\ \alpha &:= r, \end{aligned} \tag{S17}$$

where $X$ represents the probability of receptor activation, $r$ denotes the number of receptors. From this set of definitions, it follows that $Y = A$ is the number of active receptors. Note that $\alpha$ from equation (S16) is a constant value, rather than a function like the placement strategy $\phi$. Therefore, Equation S17 agrees with our local formulation of (S5), and matches the general formulation of (S2) if $\phi$ is a constant function. An important consequence of this mapping is that we can now study the quantity $I(X, Y)$ since,

$$I(X; Y) = I(f(C); A) \tag{S18}$$

$$= I(C; A), \tag{S19}$$

where the second line follows from the fact that the mutual information is invariant to invertible transformations $f$. Since most physical models of receptor activity ($f$) are strictly increasing functions of ligand count, hence invertible, theoretical properties of $I(X, Y)$ which we discuss here directly applies to many receptor models beyond what is considered in this work, such as models with signal amplification and receptor cooperativity. Note that we are leaving the probability distribution $P(X)$ unspecified, which again makes many of the following results valid for many choices of $f$.

*2.2. Relating properties of $I(X, Y)$ to receptor sensing*

Having established the relationship $I(X;Y) = I(C;A)$, we now use the scalar Poisson model to illustrate how theoretical properties of the mutual information $I(X, Y)$ agrees with our intuition of ligand sensing via receptor binding. We specialize to the case where $X$ is a non-negative random variable which is sufficient for our problem as $X$ only takes values between 0 and 1. In this setting, Theorem 2 of [8] gives the partial information gain for the scalar Poisson channel as,

$$\frac{d}{dr} I(X;Y) = E[X \log X - E[X|Y] \log E[X|Y]]. \tag{S20}$$

An immediate consequence of Equation S20 is that the mutual information $I(X, Y)$ (hence $I(C;A)$) is strictly increasing in the scaling variable (receptor number), which follows from the fact that the right side of Equation S20 is non-negative due to Jensen's inequality since $x \log x$ is a convex function. The fact that $I(X, Y)$ is strictly increasing in $r$ agrees with the intuition that increasing the number of receptors should increase the amount of information the cell can acquire about its external environment.

Observe that the right hand side of Equation S20 is exactly the minimum mean loss in estimating $X$ based on $Y$ under the loss function $l(x_1, x_2) = x_1 \log(x_1/x_2) - x_1 + x_2$. Using this fact, one can show that $I(X, Y)$ is a concave function of $r$, which again agrees with the intuition that since the total amount of information available $H(X)$ is fixed, incremental gain in information acquisition must diminish as more receptors are added. Importantly, the fact that $I(X;Y)$ is an increasing, concave function of the scaling variable $r$ holds across all models of receptor activation (Equation S17). In particularly, the concavity of $I(X;Y)$ is a general phenomena and not a result of saturation from ligand binding.

*2.3. Mapping full membrane receptor model to vector Poisson channel model*

We will now rewrite our local optimization problem of Equation S5 using the canonical vector Poisson channel model. By doing so, we will be able to use theoretical properties of the vector Poisson model to provide additional insight into the optimal solution, and expand the result beyond the specific receptor model used in the main text. By a similar argument as in the scalar Poisson case, the full membrane receptor model considered in our work (main text, Equation 3) maps exactly onto the canonical vector Poisson channel model, defined as

$$\boldsymbol{Y}|\boldsymbol{X} \sim \prod_{i=1}^{m} P(Y_i \mid \boldsymbol{X}) = \prod_{i=1}^{m} \mathrm{Pois}(Y_i \mid (\Phi \boldsymbol{X})_i) \tag{S21}$$

where the random vector $\boldsymbol{X} = (X_1, X_2, ..., X_m)$ maps to the probability of receptor activation across the $m$ discretized membrane regions, the random vector $\boldsymbol{Y} = (Y_1, Y_2, ..., Y_m)$ maps to the random vector of active receptors $\boldsymbol{A} = (A_1, A_2, ..., A_m)$, the channel matrix $\Phi \in \mathbb{R}_+^{m \times m}$ can map onto receptor placement $\boldsymbol{r} = (r_1, r_2, ..., r_m)$ such that $\Phi = \mathrm{diag}\,(\boldsymbol{r})$. This mapping represents the fact that receptors bind ligands locally and activate independently of other receptors. We introduce $\Phi$ for completeness, showing that this model can accomodate situations where there are crosstalks between channels, leading to non-zero terms in the off-diagonal. Since the equivalent of (S19) holds for the vector model, we can rewrite the optimization problem in Equation S5 as,

$$\boldsymbol{r}^* = \underset{\substack{\boldsymbol{r} \geq 0 \\ \sum_i r_i = N}}{\mathrm{argmax}}\ I(\boldsymbol{X}, \boldsymbol{Y} \mid \boldsymbol{r}). \tag{S22}$$

By working with Equation S22, we derive results that hold for many models of receptor activation, including all models where activity is a monotonic function of ligand level.

*2.4. Reformulation of receptor optimization in terms of partial information gain*

We reformulate Equation S22 in terms of the partial derivatives $\partial I(\boldsymbol{X};\boldsymbol{Y})/\partial r_i$, which provides additional insight into the optimal solution. According to the Karush-Kuhn-Tucker (KKT) conditions, the following must hold at the optimal solution $\boldsymbol{r}^*$ for $1 \leq i \leq m$ ,

$$\frac{d}{dr_i} I(\boldsymbol{X};\boldsymbol{Y} \mid \boldsymbol{r}^*) = \lambda - \mu_i, \qquad \mu_i r_i^* = 0, \tag{S23}$$

where $\mu_i \geq 0$ and $\lambda$ are the KKT multipliers. Another way to interpret the equations above is that for all channels where the optimal receptor number $r_i^*$ is non-zero, their partial derivatives $\frac{d}{dr_i} I(\boldsymbol{X};\boldsymbol{Y}) \mid_{\boldsymbol{r}^*}$ must be equal. Put another way, optimal solution occurs when incremental information gain is matched across channels. Since whenever the partial derivatives do not all agree, then one can always move receptors from the channel with a smaller partial derivative to one with higher partial derivative to achieve a higher mutual information.

*2.5. Asymptotic of the gradient of mutual information at small N*

We show that when total receptor number is low, the partial information gain depends only on the properties of $\boldsymbol{X}$, the probability of receptor activation. This result allows us to directly solve for the optimal solution at the low $N$ regime. Theorem 1 of [9] gives the gradient of mutual information between input and output of the vector Poisson channel $I(\boldsymbol{X};\boldsymbol{Y})$, with respect to the matrix $\Phi$ as,

$$\nabla_\Phi I(\boldsymbol{X};\boldsymbol{Y})_{ij} = E[X_j \log(\Phi \boldsymbol{X})_i] - E[E[X_j|\boldsymbol{Y}] \log E[(\Phi \boldsymbol{X})_i|\boldsymbol{Y}]]. \tag{S24}$$

Specializing to the setting where $\Phi$ is a diagonal matrix with $\mathrm{diag}(\Phi) = \boldsymbol{r}$, the derivative of mutual information with respect to receptor number at position $i$ is,

$$\frac{d}{dr_i} I(\boldsymbol{X};\boldsymbol{Y}) = E[X_i \log X_i] - E[E[X_i|\boldsymbol{Y}] \log E[X_i|\boldsymbol{Y}]]. \tag{S25}$$

This derivative can be interpreted as the information gain at the $i$-th channel per receptor added.

When total receptor number ($N$) is low, corresponding to all scaling variables ($\{r_i\}$) being small, we can express (S25) as a function of just the random variable $\boldsymbol{X}$. First, using Lemma 1 from [10], we have

$$\begin{aligned} r_i E[X_i|\boldsymbol{Y} = \boldsymbol{y}] &= (y_i + 1)\frac{p_{\boldsymbol{Y}}(\boldsymbol{y} + 1_i)}{p_{\boldsymbol{Y}}(\boldsymbol{y})} \\ &= r_i \frac{E\left[(X_i)^{y_i+1} e^{-r_i X_i} \prod_{m \neq i} \frac{1}{y_m!} (X_m)^{y_m} e^{-r_m X_m}\right]}{E\left[\prod_m (X_m)^{y_m} e^{-r_m X_m}\right]}. \end{aligned} \tag{S26}$$

Therefore, for $r_i > 0$, we have

$$E[X_i|\boldsymbol{Y} = \boldsymbol{y}] = \frac{E\left[(X_i)^{y_i+1} e^{-r_i X_i} \prod_{m \neq i} \frac{1}{y_m!} (X_m)^{y_m} e^{-r_m X_m}\right]}{E\left[\prod_m (X_m)^{y_m} e^{-r_m X_m}\right]}. \tag{S27}$$

Now using monotone convergence theorem, we obtain

$$\lim_{r_1,\ldots,r_m \to 0^+} E\left[X_i \mid \mathbf{Y} = \mathbf{y}\right] = \frac{E\left[(X_i)^{y_i+1} \prod_{m \neq i} \frac{1}{y_m!} (X_m)^{y_m} e^{-r_m X_m}\right]}{E\left[\prod_m (X_m)^{y_m}\right]} \tag{S28}$$

The above limit holds for any path, and also holds for all value of $\boldsymbol{y}$ including zero. Evaluating the above limit at $\boldsymbol{y} = 0$ we have

$$\lim_{r_1,\ldots,r_m \to 0^+} E[X_i|\boldsymbol{Y} = \mathbf{0}] = E[X_i]. \tag{S29}$$

Applying this limit to (S25) gives the desired result,

$$\lim_{r_1,\ldots,r_m \to 0^+} \frac{\partial}{\partial r_i} I(\boldsymbol{X};\boldsymbol{Y}) = E[X_i \log X_i] - E[X_i] \log E[X_i], \tag{S30}$$

which we denote as

$$\frac{\partial I_0}{\partial r_i} := \lim_{r_1,\ldots,r_m \to 0^+} \frac{\partial}{\partial r_i} I(\boldsymbol{X};\boldsymbol{Y}). \tag{S31}$$

In this limit, the partial derivatives are independent of receptor number. Intuitively, when receptor numbers are low, the effect of diminishing return that comes from having many receptors should be weak. Importantly, this result holds for arbitrary distribution $P(\boldsymbol{X})$, hence it holds for any environmental statistic and model of receptor activation. As we show in the next section, this fact allows us to solve for the optimal solution $\boldsymbol{r}^*$ exactly in the low $N$ limit.

#### 3. Factors affecting optimal receptor placement

##### *3.1. Total receptor number*

Optimal receptor placement can be strongly localized when receptors are limited in quantity. When receptor number is small, equation (S30) shows that the $\frac{d}{dr_i} I(\boldsymbol{X};\boldsymbol{Y})$ becomes independent of receptor number. This result implies that $I(\boldsymbol{X};\boldsymbol{Y})$ is maximized when all receptors are allocated to the channel with the largest partial derivative $\frac{d}{dr_i} I(\boldsymbol{X};\boldsymbol{Y})$, resulting in strong receptor localization. We can see this by first noting that in the limit of small $N$, equation (S30) and Taylor's theorem allows us to write the mutual information as a linear function of $\boldsymbol{r}$,

$$I(\boldsymbol{X};\boldsymbol{Y}) = \sum_{i=1}^{m} \frac{\partial I_0}{\partial r_i} r_i \tag{S32}$$

Hence, our optimization problem becomes a linear program with the following form,

$$\begin{array}{ll} \text{maximize} & \boldsymbol{a}^T \boldsymbol{r} \\ \text{subject to} & \boldsymbol{1}^T \boldsymbol{r} = r_{\text{tot}}, \quad \boldsymbol{r} \geq 0, \end{array} \tag{S33}$$

where $a_i = \partial I_0 / \partial r_i$. Suppose the $a_i$'s are sorted in increasing order (with corresponding $r_i$'s rearranged as well),

$$a_1 \leq a_2 \leq \cdots < a_k = \cdots = a_{m-1} = a_m \tag{S34}$$

Denote $a_{\max} := a_m$, we have

$$\boldsymbol{a}^T \boldsymbol{r} \leq a_{\max} (\boldsymbol{1}^T \boldsymbol{r}) = a_{\max} r_{\text{tot}} \tag{S35}$$

for all feasible $\boldsymbol{r}$, with equality if and only if

$$r_k + \cdots + r_m = r_{\text{tot}}. \tag{S36}$$

The optimal solution is then to allocate all receptors among the channels with maximal partial information gain $\partial I_0 / \partial r_i$, in any manner. This result is quite intuitive. If the channel information gains are fixed, we allocate all receptors to the channel with the highest gain. Since equation (S30) is valid for all non-negative random variable $X$, this result holds for arbitrary environmental statistics. Furthermore, since $\partial I_0 / \partial r_i = a_i \geq 0$ due to Jensen's inequality, the optimal solution remains unchanged if we replace the equality constraint by an inequality $\boldsymbol{1}^T \boldsymbol{r} \leq N$. Again, since we allow $P(\boldsymbol{X})$ to be arbitrary, this optimal solution holds for any environmental statistic and model of receptor activation. Figure S2 illustrates this result by optimizing across two Poisson channels for different number of receptors ($N$).

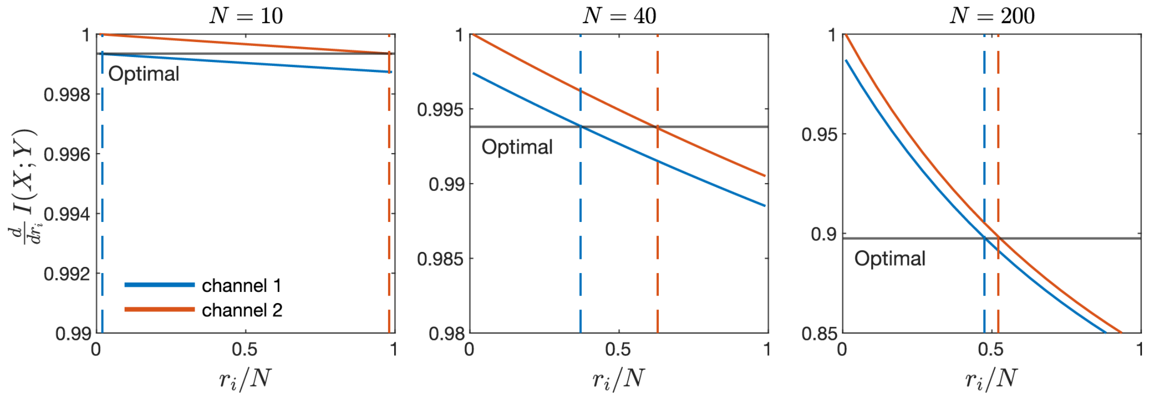


Figure S2: **Optimal receptor distribution across two Poisson channels for various values of $N$, Related to Figure 2 and Figure 7.** Given two Poisson channel with independent inputs $X_1$ and $X_2$, where $E[X_2] > E[X_1]$. Each plot corresponds to a particular choice of $N$ to be divided between the two channels. Solid curves represent the partial derivative the mutual information $I(X;Y)$ with respect to the two channel receptor number $r_1$ and $r_2$, evaluated for different receptor allocations. Dotted horizontal lines indicate the receptor distribution between the two channels that maximizes $I(X;Y)$.

In line with the KKT condition of (S23), Figure S2 shows the optimal receptor distribution across the two channels (dotted lines) occurs precisely where their partial derivatives (solid lines) are equal. Even though channel 2 (orange) experience an average ligand concentration that is only 5% higher than channel 1, the optimal solution allocates nearly all receptors (98%) to channel 2 when $N$ is small. As $N$ increases, this asymmetry of the optimal solution reduces significantly, resulting in a 5% difference in receptor number between the two channels when $N = 200$. In agreement with results we derived, strong receptor localization occurs when $N$ is small due to the fact that $\frac{d}{dr_i} I(X;Y)$ becomes nearly independent of receptor number (note the difference in y-range across the three plots in Figure S2).

##### *3.2. Absolute ligand concentration and dynamic range*

In addition to receptor number, environmental factors can strongly influence receptor placement. Intuitively, one would expect the larger the difference between two channels' input ligand concentration, the larger the asymmetry should be in their receptor allocation. Figure S3A confirms this intuition, showing that as the relative difference in average ligand concentration sensed between two channels increase, receptor distribution between the two channels becomes more asymmetric. Figure S3A also suggests two additional features of the optimal strategy that are less intuitive,

1. As ligand concentration increases, optimal strategy switches from allocating more to allocating less receptors to region of higher ligand concentration
2. When ligand concentration are either high or low, optimal receptor placement can become highly localized, concentrating most receptors to a few channels

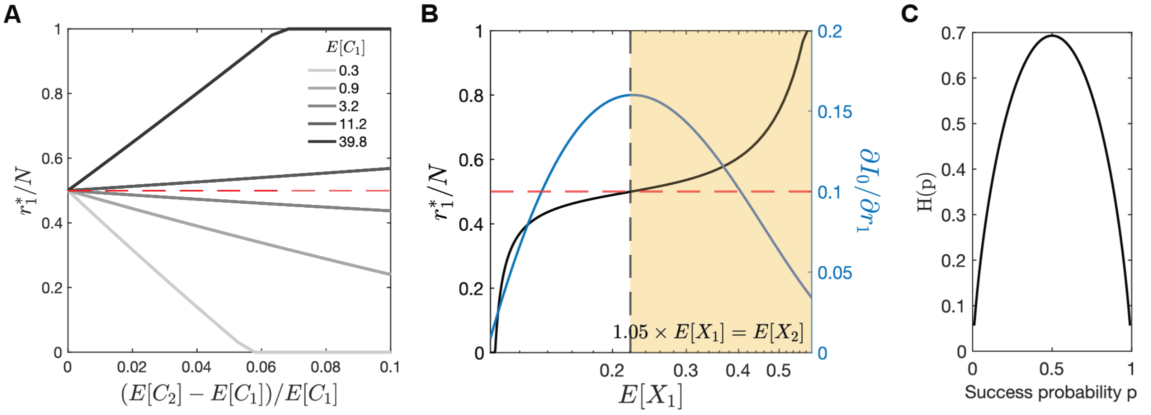


Figure S3: **Optimal receptor distribution across two Poisson channels for different absolute ligand concentration and relative difference in concentration, Related to Figure 2.** (A) Each curve represents the optimal receptor proportion $r_1^*/N$ for channel 1 for increasing levels of ligand input for channel 2 while keeping $E[C_1]$ fixed. (B) While keeping the relative difference between $E[C_1]$ and $E[C_2]$ fixed at 5%, black curve shows the optimal receptor proportion in channel 1 for different level of $E[C_1]$. Blue curve shows the approximation of the partial derivative shown in equation (S30). Black dotted line indicates peak of blue curve, red dotted line indicated $r_1^*/N = 0.5$. $K_d = 40nM$. (C) binary entropy function ($H(p) = -p \log p - (1-p) \log (1-p)$) in nats

The first feature can be seen by observing the fact that as $E[C_1]$ increases in Figure S3A, the optimal receptor distribution $r_1^*/N$ (gray to black) changes from being below 0.5 to above 0.5, even though $E[C_2] > E[C_1]$ for all cases plotted. The second case can be seen by observing the slope of the graphs. As $E[C_1]$ becomes either high (black) or low (gray), the optimal receptor distribution becomes more sensitive to $(E[C_2] - E[C_1])/E[C_1]$, the relative difference in average input level. For a minor difference in concentration of $< 5\%$, nearly all receptors become allocated to one of the two channels

Both of these observations are indeed general features of the optimal strategy and we can explain both using $\partial I_0/\partial r_i$ defined in (S30). We can gain further intuition of both features from the shape of the binary entropy function (Figure S3C).

1. Figure S3B shows that when $E[X_1]$ is low, $\partial I_0/\partial r_1$ is an increasing function in $E[X_1]$, suggesting that more receptors should be allocated to channels with higher probability of receptor activation (i.e. ligand concentration). However, this monotonicity switches as $E[X_1]$ increases, with $\partial I_0/\partial r_1$ becoming a decreasing function in $E[X_1]$. The point at which monotonicity of $\partial I_0/\partial r_1$ switches (dashed black line) matches precisely with when the optimal strategy switches from allocating more receptors to region of higher probability of receptor activation to region of lower probability, as shown by the solid black curve passing the dashed red line. Thus feature #1 can be fully explained by the gradient of mutual information. This switch in strategy is intuitive when we consider the binary entropy function. Recall that $X_i$ maps to the probability of receptor activation in the $i$-th channel, so it shares a similar interpretation as the success probability $p$ of the binary entropy function. The entropy function $H(p)$ is maximized when the success probability (analogously receptor activation) is neither high nor low (Figure S3C). Thus it can be less useful to place more receptors at regions of higher ligand concentration, since those receptors will simply stay activated, being uninformative of the input.
2. Figure S3B shows that the slope of $\partial I_0/\partial r_1$ is maximized when $E[X_1]$ is either low or high. The larger the difference in the partial derivatives between two channels, the more receptors will need to be allocated before their partial derivative agree, a necessary condition for achieving optimality according to (S23). Therefore, a small relative difference in input concentration between channels can lead to large difference in information gain per receptor, when absolute ligand concentration is either high or low (relative to $K_d$), leading to strong localization of receptors. This behavior can also be explained using the binary entropy function, specifically the fact that the rate of change in entropy is maximized at low and high success probability (Figure S3C). Analogously,

placing receptors in regions where likelihood of activation is 0 or 1 is useless from an information perspective (zero entropy/uncertainty in output), so all receptors should be allocated to a region with non-zero entropy, no matter how small the difference in likelihood of activation (thus ligand concentration) is.

#### 4. Modeling chemical microenvironment

##### *4.1. Soil chemical microenvironment*

In soil, free-living unicellular eukaryotes can sense and respond to signaling ligands secreted by soil bacteria. We follow mathematical models described in [4] and [3], modeling the spatial distribution of ligands in two steps: 1) model the spatial distribution of bacteria in soil, and 2) model each bacteria as an independent point sources of ligands.

*Modeling bacteria distribution in soil.* We follow the procedure outlined Raynaud and Nunan (2014) [4], which allows us to generate realistic bacterial distributions found in soil. This procedure involves sampling from a spatial statistical model, based on Log Gaussian Cox Processes (LGCP) fitted to image data of observed bacterial distribution in soil. In a LGCP, the observed number of bacteria per unit area is modeled as a Poisson process in which the rate parameter is treated as being the exponential of a Gaussian process. Specifically, we consider Gaussian processes with an exponential covariance function,

$$C(r) = \sigma^2_{\text{bacteria}} e^{-r/\beta}, \tag{S37}$$

so the Gaussian process (and the LGCP) is fully determined by three parameters, its mean ($\mu$), variance ($\sigma^2_{\text{bacteria}}$), and scale ($\beta$). In the limit as $\sigma^2_{\text{bacteria}} \to 0$, we obtain a homogeneous Poisson process. The average intensity of a LGCP (number of bacteria per unit area) is given by,

$$\lambda = e^{\mu + \sigma^2_{\text{bacteria}}/2} \tag{S38}$$

We used parameters reported in [4], with $\mu = -7.52$, $\sigma^2_{\text{bacteria}} = 1.9$, and $\beta = 25$, to simulate a bacterial density of approximately $10^9$ cells/g on a $1000 \times 3000\mu\text{m}^2$ rectangular domain (containing approx. 4000 cells). These are the default parameters unless otherwise stated in the main text. We used R 3.6.1 with packages *spatstat* [11] and *RandomFields* [12] to generate all bacteria distributions.

*Modeling chemical distribution given bacteria distribution..* Given a spatial distribution of bacteria, we model the distribution of secreted molecules using standard reaction-diffusion models [3]. Specifically, such models treat each bacteria as a static, independent sources, producing ligands with rate $\alpha$ that diffuse ($D$) and degrade ($\gamma$). The resulting ligand concentration field is then the solution of the following partial-differential equation (PDE) ,

$$\frac{\partial c(x,t)}{\partial t} = \alpha|_{\text{bacteria}} + D\Delta c - \gamma c, \tag{S39}$$

Rather than approximating each parameter of (S39), we model the ligand distribution produced by a bacteria using a 2D Gaussian density profile, and directly fit the Gaussian profile to empirical measurements. The concentration $c$ at a given position $x$ in the domain is then the sum over all such Gaussian profiles evaluated at $x$, which can be expressed mathematically as

$$c(x) = \sum_{q \in \mathcal{U}} \frac{C}{\sqrt{2\pi s^2}} \exp\left\{ -\frac{||x-q||^2}{2s^2} \right\} \tag{S40}$$

where $\mathcal{U}$ represents the set of bacterial positions generated using the LGCP model. $C$ represents the total concentration of each Gaussian profile, and $s$ determines the width of the profile, both of which are assumed to be uniform across all bacteria. We extract both parameters based on a geostatistical block kriging analysis of the spatial distribution of AHL in soil ($C$ chosen such that mean concentration (across the entire spatial domain) is approximately 0.6 nM, $s = 9\,\mu\text{m}$) [13, 14, 15, 16].

*4.2. Tissue chemical microenvironment*

We follow mathematical models of ligand distribution in tissue outlined in [6, 5], simulating a tissue environment using a PDE model that incorporates four transport mechanisms: (1) free diffusion, (2) ECM binding, (3) fluid advection, (4) cellular uptake. The spatial domain is a rectangle of size $300\,\mu\text{m} \times 900\,\mu\text{m}$. We model ligands being supplied through fluid flows from the left boundary of the domain, and penetrate the interstitial space between immobilized cells. Soluble ligands are then transported by diffusion and fluid flow, and become immobilized upon binding to an extracellular matrix (ECM) made up of networks of interconnected fibers containing ligand binding sites. We explicitly represent both ECM-bound ($c_b$) and soluble forms of the ligand ($c_s$), so that the the total ligand concentration $c(x,t)$ at position $x$ and time $t$ is equal to,

$$c(x,t) = c_s(x,t) + c_b(x,t). \tag{S41}$$

Mathematically, we can describe the dynamics of the soluble fraction $c_s(x,t)$ as follows,

$$\frac{\partial c_s}{\partial t} = \kappa|_{\text{boundary}} - u(x,t)\cdot\nabla c_s + D\Delta c_s - \beta_c c_s|_{\text{cells}} - k_{\text{ECM}}(e(x) - c_b)c_s - \gamma_s c_s. \tag{S42}$$

1. The first term, $\kappa$, represents production/release of molecule at the left boundary.
2. The second term represents fluid transport, where $u(x,t)$ is the velocity field of the interstitial fluid with input flow speed $u^{\text{in}}$ at the left boundary. We impose zero-velocity condition on the top and bottom boundary.
3. The third term represents diffusion with $D$ as the ligand diffusion coefficient.
4. The fourth term represents cellular uptake with rate $\beta_c$, a process that only occurs near immobilized cells distributed across the domain.
5. The 5th term represents ECM binding. The concentration of ECM binding site $e(x)$ at position $x$ is generated using a minimal model of ECM protein distribution (see paragraph on "Generating ECM fiber network"). Binding occur with rate proportional to $e(x)-c_b(x,t)$, the level of available ECM binding site. Since the on-rate of ECM binding is much larger than the off-rate, we assume the off-rate to be zero.
6. The last term represents enzymatic degradation of ligand.

The dynamics of ECM-bound fraction $c_b(x,t)$ is much simpler, involving a term corresponding to ECM binding, a degradation term due to enzymatic decay .

$$\frac{\partial c_b}{\partial t} = k_{\text{ECM}}(e(x) - c_b)c_s - \gamma_b c_b. \tag{S43}$$

To generate a ligand concentration field $c$, we take $\kappa$ to be non-zero for a brief period of time, representing a bolus of ligand released. Then, we simulate the combined dynamics of bound and soluble fractions for sufficiently long until the ligand distribution $c(x,t)$ is relatively stable. In practice, we observe that $c \approx c_b$ after a sufficiently long period of time, since the soluble fraction quickly become insignificant due to fluid flow. The resulting concentration field represents an interstitial gradient. The average concentration is set by setting the release rate $\kappa$ such that the concentration of the soluble fraction $c_s$ matches measured chemokine concentration found in interstitial fluids (1 pM−10 pM) [17, 18].

*Generating ECM fiber network.* To generate a distribution of ECM binding sites $e(x)$, we use a minimal computation model of fiber network [19, 20, 21]. The model generates ECM fibers represented by line segments, which could represent fibronectin, collagen, laminin, or other fibrous matrix components. To position each fiber, one end of each segment is randomly positioned following a uniform distribution within the domain. The other end's position is determined by picking an angle, uniformly from $[0, 2\pi)$, and length sampled from a normal distribution with mean 75 µm and standard deviation of 5 µm (as measured for collagen by Friedl et al [22]). In total, 4050 fibers were placed in the domain. For the PDE simulation, the generated network is discretized by counting the number of fibrous proteins around each node in the simulation lattice. The density of fiber within each node is then converted to a concentration value representing the level of ECM binding sites, resulting in an average concentration of ECM binding site of 520 nM.

*4.3. Simple chemical gradient*

One of the simplest model of chemical gradient can be described by the following PDE,

$$\frac{\partial c(x,t)}{\partial t} = \alpha\delta(x_0) + D\Delta c - \gamma c, \tag{S44}$$

where ligands are produced at rate $\alpha$ from a localized source at $x_0$, diffuses with diffusivity $D$ and undergoes first order degradation with rate $\gamma$. The steady-state solution of (S44) is a single exponential gradient,

$$c(x) = C_0 \exp(-x/\lambda), \tag{S45}$$

where the ligand concentration $c(x)$ only depends on distance $x$ from the source, the concentration at the source boundary $C_0 = \alpha/(2\sqrt{D/\gamma})$ and the decay length $\lambda = \sqrt{D/\gamma}$. By taking the source location $x_0$ to be the entire left boundary of the spatial domain, the stimulated interstitial gradient is well-described by the exponential model. Specifically, by first averaging the interstitial gradient (along the axis parallel to the ligand source) and fitting the resulting 1-D profile to Equation S45 using Matlab's fit function, we obtain an excellent fit with correlation coefficient $R^2 = 0.98$. This fitted exponential profile is the simple, monotonic gradient used in the paper.

| Environment class | Parameter | Symbol | Value | Ref |
|---|---|---|---|---|
| Soil | Domain size | — | $1000\,\mu m \times 3000\,\mu m$ | — |
| | mean | $\mu$ | $-7.52$ | [4] |
| | variance | $\sigma^2_{\text{bacteria}}$ | 1.9 | [4] |
| | scale | $\beta$ | 25 | [4] |
| | Concentration (per bacteria) | $C$ | 114 nM | [13, 15, 16] |
| | Spread of ligand (per bacteria) | $s$ | 9 µm | [14] |
| Tissue | Domain size | — | $300\,\mu m \times 900\,\mu m$ | — |
| | Diffusion coefficient | $D$ | $45\,\mu m^2\,s^{-1}$ | [23] |
| | Cellular uptake rate | $\beta_c$ | $10^{-2}\,s^{-1}$ | [6] |
| | Cellular uptake distance | — | 1.15× radius | [6] |
| | Fluid viscosity | $\mu$ | $2.5\,\mu g\,\mu m^{-1}\,s^{-1}$ | [6] |
| | Spatial discretization | $\Delta x$ | 2 µm | — |
| | Time step | $\Delta t$ | 0.0178 s | — |
| | Interstitial fluid input flow | $u^{\text{in}}$ | $0.1\,\mu m\,s^{-1} - 2\,\mu m\,s^{-1}$ | [24] |
| | Average [ECM binding site] | — | 520 nM | [25] |
| | ECM binding rate | $k_{\text{ECM}}$ | $9.3 \times 10^{-5}\,nM^{-1}\,s^{-1}$ | [26] |
| | Production/release rate | $\kappa$ | $7\,nM\,s^{-1}$ | [17] |
| | Number of cells | — | $3 \times 8$ cells | — |
| | Soluble ligand degradation | $\gamma_s$ | $1 \times 10^{-3}\,s^{-1}$ | [5, 25] |
| | Bound ligand degradation | $\gamma_b$ | $1 \times 10^{-5}\,s^{-1}$ | [5] |
| | Mean ECM fiber length | — | 75 µm | [20] |
| | Variance in fiber length | — | 5 µm | [20] |
| | Total number of ECM fibers | — | 4050 | [20] |
| Simple gradient | Max concentration | $C_0$ | 0.7 nM | – |
| | Decay length | $\lambda$ | 60 µm | – |

Table S1: **Parameters used for modeling all three environment classes: soil, tissue, and simple gradient, Related to Figure 2A.** Tissue simulation code was adopted from [6]. Parameters for simple gradient obtained from fitting an exponential function to the spatially-averaged profile of the tissue gradient

#### 5. Incorporate cost for receptor redistribution using the Wasserstein distance

In a dynamically changing environment, receptor should redistribute in an efficient manner in order to maximize information acquisition. We extended our optimization problem of equation (S5)

to incorporate a "cost" for changing receptor location. For a cell sensing a sequence of ligand profiles $\{\boldsymbol{c}_t\}_{t=1}^T$ over time, the optimal receptor placement $\boldsymbol{r}_t^*$ for $\boldsymbol{c}_t$ now depends additionally on $\boldsymbol{r}_{t-1}^*$, the receptor placement for the previous ligand profile,

$$\boldsymbol{r}_t^* = \underset{\substack{\boldsymbol{r}\geq 0 \\ \sum_i r_i = N}}{\operatorname{argmax}} \; I\left(\hat{\boldsymbol{c}}_t; \hat{\boldsymbol{a}} \mid \boldsymbol{r}\right) - \gamma W_1\left(\boldsymbol{r}_{t-1}^*, \boldsymbol{r}\right). \tag{S46}$$

Here, we model the cost for redistributing receptors using the Wasserstein-1 ($W_1$) distance. For completeness, we first introduce the formal definition of the $W_1$ distance before returning to a much simpler form that applies to our problem. Let $X \sim P$ and $Y \sim Q$ represent two random variables defined over $M \subset \mathbb{R}^d$. Further, let $\mathcal{J}(P,Q)$ denote all joint distributions $J$ for $(X,Y)$ that have marginal $P$ and $Q$. The $W_1$ distance between $P$ and $Q$ is,

$$W_1(P,Q) = \inf_{J\in\mathcal{J}(P,Q)} \int_{M\times M} \|x-y\|_1 dJ(x,y). \tag{S47}$$

One way to understand the above definition is to consider different ways of transporting a distribution of mass $P(x)$ to a different distribution $Q(x)$. Given some cost function associated with each unit of mass transported, the $W_1$ distance is the minimum transport cost achievable. In this way, the $W_1$ distance assumes that the transformation from $P$ to $Q$ occurs in an optimal manner. Note that this distance function is non-negative and symmetric, and does not require $P$ and $Q$ to be probability distributions, it applies whenever the total mass is preserved between $P$ and $Q$.

Although equation (S47) is difficult to compute in general, it has a closed form for the special case of $d = 1$ which is the case we are considering. Instead of using the canonical form of the $W_1$ distance in 1-D, we need to use a generalized form that applies to distributions on a circle [27]. For two receptor distributions on the 1-D surface of a 2-D cell, represented as non-negative vectors $\boldsymbol{a}$ and $\boldsymbol{b}$ of length $m$, the $W_1$ distance takes on the form,

$$W_1(\boldsymbol{a},\boldsymbol{b}) = \sum_{i=1}^{m} |\phi_i - \mu|, \tag{S48}$$

where $\phi_i = \sum_{j=1}^{i}\left(\frac{a_j}{\|a\|_1} - \frac{b_j}{\|b\|_1}\right)$ and $\mu$ is the median of the set of values $\{\phi_i, 1 \leq i \leq m\}$. We derive the gradient of equation (S48) as,

$$\frac{\partial}{\partial a_k} W_1(\boldsymbol{a},\boldsymbol{b}) = \sum_{i=1}^{m} \operatorname{sgn}(\phi_i - \mu) \sum_{j=1}^{i}\left(\delta_{jk} - \frac{a_j}{\|a\|_1}\right). \tag{S49}$$

We perform optimization with this gradient using the fmincon function (with sqp algorithm) in Matlab.

#### 6. Numerical simulation of receptor feedback scheme

In our feedback scheme, receptor $r(x,t)$ is modeled by considering three redistribution mechanisms: (1) lateral diffusion of $r$ along the plasma membrane ($D\nabla^2_{\mathrm{memb}} r$), (2) endocytosis of $r$ along the plasma membrane ($k_{\mathrm{off}}\, r$), (3) incorporation of cytoplasmic pool of receptors, $R_{\mathrm{cyto}}$, to the membrane at rate proportional to local receptor activity ($hAR_{\mathrm{cyto}}$). $A(x,t)$ is a random variable that denotes receptor activity along the cell membrane, and is a function of local receptor number. Then, the equation describing the distribution of $r$ across the cell membrane can be expressed mathematically as,

$$\frac{\partial r(x,t)}{\partial t} = D\nabla^2_{\mathrm{memb}} r - k_{\mathrm{off}}\, r + hAR_{\mathrm{cyto}}, \tag{S50}$$

where the total number of receptors $r_{\mathrm{tot}} = \int_{\mathrm{memb}} r + R_{\mathrm{cyto}}$ is fixed. We simulate receptor distribution by treating the cell membrane as a 1D space and the cytosol as a single, homogeneous compartment.

This simplification allows us to simulate our PDE using the Crank-Nicolson method in one spatial dimension. Given space and time units $\Delta x$ and $\Delta t$, respectively, the Crank-Nicolson method with $R_i^j := r(i\Delta x, j\Delta t)$ and $A_i^j := A(i\Delta x, j\Delta t)$ is given by the difference scheme

$$\begin{aligned}\frac{R_i^{j+1} - R_i^j}{\Delta t} = \frac{D}{2\Delta x^2}\Big(R_{i+1}^j - 2R_i^j + R_{i-1}^j + R_{i+1}^{j+1} - 2R_i^{j+1} + R_{i-1}^{j+1}\Big) \\ - \frac{k_{\text{off}}}{2}\Big(R_i^j + R_i^{j+1}\Big) + \frac{hA_i^j}{2}\Big(R_{\text{cyto}}^j + R_{\text{cyto}}^{j+1}\Big)\end{aligned} \tag{S51}$$

where, $i = 1, 2, 3, ...m$, representing $m$ discrete membrane compartments and $R_{\text{cyto}}^j$ represents the additional cytosol compartment. Since the membrane is represented by a circle, we have the following pair of conditions,

$$R_0^j = R_m^j, \;\; R_{m+1}^j = R_1^j. \tag{S52}$$

Lastly, total receptor number across all compartments is conserved,

$$\sum_{i=1}^m R_i^j + R_{\text{cyto}}^j = \sum_{i=1}^m R_i^{j+1} + R_{\text{cyto}}^{j+1}. \tag{S53}$$

Now, we can combined (S51)-(S53) and rewrite everything in vector form. First, let

$$\alpha := \frac{D}{2\Delta x^2}, \;\; \beta := \frac{k_{\text{off}}}{2}, \;\; \kappa_i^j := \frac{hA_i^j}{2},$$

and rewrite equation (S51) as,

$$\begin{aligned}\frac{R_i^{j+1}}{\Delta t} - \alpha\Big(R_{i+1}^{j+1} - 2R_i^{j+1} + R_{i-1}^{j+1}\Big) + \beta R_i^{j+1} - \kappa_i^{j+1} R_{\text{cyto}}^{j+1} = \frac{R_i^j}{\Delta t} + \alpha\Big(R_{i+1}^j - 2R_i^j + R_{i-1}^j\Big) \\ - \beta R_i^j + \kappa_i^j R_{\text{cyto}}^j\end{aligned} \tag{S54}$$

and define $U^j$ to be the $(m+1)$-dimensional vector with components $R_i^j$ for $i = 1, 2, 3, ...m$ and $U_{m+1}^j = R_{\text{cyto}}^j$. The difference scheme is given in the vector form

$$PU^{j+1} = QU^j. \tag{S55}$$

where,

$$P = \begin{bmatrix} \frac{1}{\Delta t} + 2\alpha + \beta & -\alpha & 0 & \cdots & 0 & -\alpha & -\kappa_1^{j+1} \\ -\alpha & \frac{1}{\Delta t} + 2\alpha + \beta & -\alpha & 0 & \cdots & 0 & -\kappa_2^{j+1} \\ 0 & \ddots & \ddots & \ddots & & & \vdots \\ \vdots & & \ddots & \ddots & \ddots & & \\ 0 & \cdots & 0 & -\alpha & \frac{1}{\Delta t} + 2\alpha + \beta & -\alpha & -\kappa_{m-1}^{j+1} \\ -\alpha & 0 & \cdots\; 0 & & -\alpha & \frac{1}{\Delta t} + 2\alpha + \beta & -\kappa_m^{j+1} \\ 1 & 1 & & \cdots & & 1 & 1 \end{bmatrix} \tag{S56}$$

$$Q = \begin{bmatrix} \frac{1}{\Delta t} - 2\alpha - \beta & \alpha & 0 & \cdots & 0 & \alpha & \kappa_1^j \\ -\alpha & \frac{1}{\Delta t} - 2\alpha - \beta & \alpha & 0 & \cdots & 0 & \kappa_2^j \\ 0 & \ddots & \ddots & \ddots & & & \vdots \\ \vdots & & \ddots & \ddots & \ddots & & \\ 0 & \cdots & 0 & \alpha & \frac{1}{\Delta t} - 2\alpha - \beta & \alpha & \kappa_{m-1}^j \\ \alpha & 0 & \cdots\; 0 & & \alpha & \frac{1}{\Delta t} - 2\alpha - \beta & \kappa_m^j \\ 1 & 1 & & \cdots & & 1 & 1 \end{bmatrix} \tag{S57}$$

Because $A$ is invertible, the Crank-Nicolson scheme reduces to the iterative process

$$U^{j+1} = P^{-1}QU^j. \tag{S58}$$

The entire evolution of $r$ can be solved where at each time step, we update receptor activity $A_i^j$ across all membrane position $i$ according to the random process described by equation (S3), (S4), followed by solving equation (S58) for $U^{j+1}$.

| Parameter | Symbol | Value | Ref. |
|---|---|---|---|
| Membrane diffusion coefficient | $D$ | $10^{-2}\,\mu m^2\,s^{-1}$ | [28, 29, 30] |
| Receptor endocytosis rate | $k_{\text{off}}$ | $0.06\,s^{-1} - 0.18\,s^{-1}$ | [31] |
| Feedback constant | $h$ | $2\times10^{-3}\,s^{-1} - 4\times10^{-3}\,s^{-1}$ | [31] |
| Average receptor feedback rate | $\langle hA_i\rangle_i$ | $10^{-3}\,s^{-1} - 2\times10^{-3}\,s^{-1}$ | [31] |
| Receptor dissociation constant | $K_d$ | $40\,\mu M$ | — |
| Cell radius | | $10\,\mu m$ | — |
| Basal receptor activity | $\alpha$ | 0.1 | — |
| Spatial discretization | $\Delta x$ | $0.6283\,\mu m$ | — |
| Time discretization | $\Delta t$ | 1 s | — |
| Total receptor | $r_{\text{tot}}$ | 1000 | — |
| Number of membrane bins | $m$ | 100 | — |

Table S2: **Parameter values for feedback scheme simulation, Related to Figure 5.** The average rate of receptor incorporation, $\langle hA_i\rangle_i$, depends on receptor activity which changes as the cell moves in a heterogeneous environment. Thus, the range shown represents the time-averaged value for a cell moving through simulated tissue environments. The value of $h$ was chosen to achieve a physiologically relevant range for $\langle hA_i\rangle_i$.

We set the value of the feedback constant $h$ using empirical measurements from Marco et al. (2007) [31]. In Figure 3M of Marco et al., the authors report a quartile box plot showing estimated values for a parameter they call h (which we will refer to as $\bar{h}$), with a mean estimate of around $1.6\times10^{-3}\,s^{-1}$. Note $\bar{h}$ is equivalent in meaning as our $hA_i$. However, since $hA_i$ will be different across different membrane bins and across time, we simulate the feedback scheme for a cell in a given environment and set the value of h such that the mean rate $\langle hA_i\rangle$ (averaged across membrane and time) is approximately equal to the mean estimate of $1.6\times10^{-3}\,s^{-1}$ reported by Marco et al.. The value $\bar{h}$ reported by Marco et al. corresponds specifically to the transport rate of the Cdc42 to the membrane. The parameter value was obtained by analyzing fluorescence recovery of GFP-Cdc42 in membrane regions bleached with a laser pulse. Although the measured value corresponds to Cdc42, it has been used to model the effective exocytosis rate for receptors shown to undergo activity-dependent localization, showing good agreement with empirical data [32]. Similar values around $10^{-3}\,s^{-1}$-$2\times10^{-3}\,s^{-1}$ have been measured for the recycling rate of a wide range of GPCRs [33, 34, 35, 36].

#### 7. Numerical simulation of cell navigating in interstitial chemical gradients

*Chemotaxis algorithm*

At $t = 0$, initialize a cell at position $p_0 \in \Omega \subset \mathbb{R}^2$.

At each subsequent time step $t = t + \Delta t$ with the cell at position $p_t \in \Omega$:

1. Compute mean ligand profile $\boldsymbol{c} \in \mathbb{R}^m$ at the cell's current position.
2. Independently sample $n$ ligand profiles $\{\boldsymbol{C}^{(i)}\}_{i=1}^n$ where each element $C_j$ is distributed as a Poisson random variable with mean equal to $c_j$ ($n = 30$ used in main text, refer to Figure S9 for other values of $n$).
3. For each ligand profile $\boldsymbol{C}^{(i)}$ sampled, sample a corresponding receptor activity profiles $\boldsymbol{A}^{(i)}$,

$$\boldsymbol{A}^{(i)}|\boldsymbol{C}^{(i)} \sim \prod_{j=1}^{m} \text{Pois}(\lambda_j), \text{ where } \lambda_j = r_j\left(\frac{C_j^{(i)}}{C_j^{(i)} + K_d} + \alpha\frac{K_d}{C_j^{(i)} + K_d}\right). \tag{S59}$$

4. Compute average receptor activity $\bar{\boldsymbol{A}} = \frac{1}{n}\sum_{i=1}^{n} \boldsymbol{A}^{(i)}$
5. Compute the an estimator of gradient direction $\hat{\theta}$ using one of three approaches
   - Optimal decoder + noise [37]: $\hat{\theta} = \arctan\left(\frac{\sin(\boldsymbol{\phi})^T \bar{\boldsymbol{A}}}{\cos(\boldsymbol{\phi})^T \bar{\boldsymbol{A}}}\right) + \mathcal{N}(0, 0.1)$, where $\phi_i = 2\pi i/m$, $i = 1, ..., m$ corresponds to the angle where $A_i$ is measured on the cell surface.
   - Random: $\hat{\theta}$ is sampled uniformly from the set of $m$ angles/directions $\{\phi_i\}_{i=1}^{m}$
   - Maximal increase: $\hat{\theta} = \phi_{i^*}$ where

$$i^* = \underset{1 \le i \le m}{\operatorname{argmax}} f(i) \quad \text{and} \quad f(i) = \begin{cases} \bar{A}_i - \bar{A}_{i+m/2} & i \le m/2 \\ \bar{A}_i - \bar{A}_{i-m/2} & i > m/2 \end{cases}$$

   This decoder selects the direction of maximum change in receptor activity across the cell surface.

   (*) In addition to the three decoders above, we consider the possibility of temporal averaging, where $\bar{\boldsymbol{A}}$ is a running mean over the past 5 minutes of receptor activity profiles (a total of $300 \times 30 = 9000$ sample profiles). This running average is decoded with the optimal decoder + noise as described above.
6. Set new cell position $p_{t+\Delta t} = p_t + s\Delta t[\cos(\hat{\theta}), \sin(\hat{\theta})]$, with speed $s = 2\,\mu\text{m}\,\text{min}^{-1}$, $\Delta t = 1\,\text{s}$.
7. Repeat from step 1.

*Tissue gradient simulation for cell navigation*

*Localization task.* In addition to using the same tissue environment as the rest of the paper, we simulated additional tissue gradients using the same set of parameters but different (randomly generated) ECM fiber networks. This results in tissue gradients that have the same macroscopic features but different patterns of microscopic fluctuations.

*Retention task.* This task was motivated by the precision with which growth cones can retain themselves within specific regions of gradients of axon guidance cues. For this task, we used an ellipse-shaped cell with semi-major axis = 5 µm, semi-minor axis = 2 µm to mimic the shape of a navigating growth cone. In-vivo observations of axon guidance cue gradients show very short decay length [38, 39], so we adjust several parameters to generate interstitial gradients with matching decay length. Below are the new parameter values that differ from Table S1,

| Parameter | Symbol | Value |
|---|---|---|
| Diffusion coefficient | $D$ | $5\,\mu\text{m}^2\,\text{s}^{-1}$ |
| Interstitial fluid flow speed | $u^{\text{in}}$ | $0.3\,\mu\text{m}\,\text{s}^{-1}$ |
| Production/release rate | $\kappa$ | $20\,\text{nM}\,\text{s}^{-1}$ |
| Soluble ligand degradation | $\gamma_s$ | $3 \times 10^{-1}\,\text{s}^{-1}$ |
| Bound ligand degradation | $\gamma_b$ | $3 \times 10^{-3}\,\text{s}^{-1}$ |

Table S3: **Parameter values used for tissue gradient generated for retention task that differs from the values in Table S1, Related to Figure 6.**

#### 8. Data on receptor placement, surface expression level, and binding affinity

Empirical measurements of receptor cell surface expression level and binding affinity can be highly variable, depending on how the affinity was measured and the particular cell type used. The data shown below simply represent a subset of values reported in literature.

| Cell type | Receptor/Ligand | $K_d$ (nM) | $N$ (cell surface) | Dynamic localization |
|---|---|---|---|---|
| T-cell | CCR2/CCL2 | 1.53 [40] | 1100 [40] | Yes [41] |
| | CXCR4/CXCL12 | 5 [42] | 1572 (834-1961) [43] | Yes [44, 45] |
| | CCR5/CCL5 | 4 [42] | 593 (167-1006) [43] | Yes[41, 46] |
| | IL-2R/IL-2 | $1.3 \times 10^{-2}$ [47] | 1000 [48] | No [41] |
| | TNFR-1/TNF | $1.9 \times 10^{-2}$ [49] | 1000 [50, 51] | No[41] |
| | TGF$\beta$R-2 | $5 \times 10^{-2}$ [52] | 10000 [53] | No [41] |
| Neutrophile | CR3/C3bi | 12.5 [54] | 40000 [55] | No [56] |
| | C5aR/C5a | 2 [57] | 50000 [57] | No [58] |
| Neuron | GABA$_A R$/GABA | 12 [59] | 200 [60] | Yes [61] |
| | Robo1/Slit | $235 \pm 165$ [62] | 22300* [63] | Yes [64] |
| | PlxnA1 | NA | NA | Yes [64] |
| Fibroblast | LPA-2/LPA | NA | NA | Yes [65] |
| Mesenchymal stem cell | CCR2/CCL2 | NA | NA | Yes [66] |

Table S4: **Receptor data used for Figure 7 of main text, Related to Figure 7.**. Receptors with known spatial distribution across cell surface, dynamic localization refers to receptors that dynamically localize towards region of highest ligand concentration, no dynamic localization refers to receptors that remain uniformly distributed even when ligands are spatially non-uniform. Only receptors with all parameter values known are included in Figure 7. (*) receptor expression level data for Robo1 was taken from a non-neuronal cancer cell line.

#### 9. Extension of information theoretic framework to study other spatial sensing strategies such as modulation of cell shape

We briefly illustrate an extension of our framework to study how cell shape can be tuned to improve cell sensing and navigation. Recall the generalized model of receptor activation from the Discussion,

$$\mathbb{E}[A_i \mid c_i] = f(\boldsymbol{\theta}_i)\Big(\frac{c_i}{c_i + K_d} + \alpha\frac{K_d}{c_i + K_d}\Big), \tag{S60}$$

where $f$ is an unspecified function of an arbitrary set of variables $\boldsymbol{\theta}$, representing the "effective" number of receptors at position $i$.

Recent work has shown that given uniform membrane receptors sensing a uniform ligand field, membrane regions of higher curvature can exhibit higher receptor activity, due to higher local volume-to-surface ratio [67]. Suppose we are interested in tuning membrane shape/curvature as a way to maximize information acquisition by cells. Assuming a constant, linear relationship between curvature at the i-th membrane position $\beta_i$ and "effective" receptor number $f$, and that receptors are uniformly distributed, then we have

$$\mathbb{E}[A_i \mid c_i] = \alpha\frac{r_{\text{tot}}}{N}\beta_i\Big(\frac{c_i}{c_i + K_d} + \alpha\frac{K_d}{c_i + K_d}\Big), \tag{S61}$$

where $\alpha$ is a proportionality constant and $N$ is the number of membrane bins. This model is identical to our receptor model (S15) up to a constant factor. Furthermore, if we assume a fixed total membrane area, the resulting optimization problem is nearly identical with (S5), where total membrane area now play a similar role as total receptor number, and $\beta_i$ takes the place of $r_i$. Therefore, we expect general features of the optimal cell shape to match that of the optimal receptor placement. Namely, cells can maximize information acquisition by increasing membrane curvature at regions of high ligand concentration, by making narrow protrusions. One can derive a more accurate solution by considering a detailed model of the relationship between curvature and receptor activity outlined in [67].

By extension, a strategy to dynamically form narrow membrane protrusions at regions of high ligand concentration, without explicitly tuning receptor positions, should in principle boost navigation efficiency in a manner similar to the receptor feedback scheme we proposed, as the two strategies have qualitatively similar effects on the spatial distribution of receptor activity. Recent works show that indeed a feedback circuit that produces dynamic, narrow membrane protrusions is crucial for neutrophil navigation. Cells that cannot form narrow protrusions can still move, but exhibit profoundly defective chemotaxis [68].

### Supplemental Figures

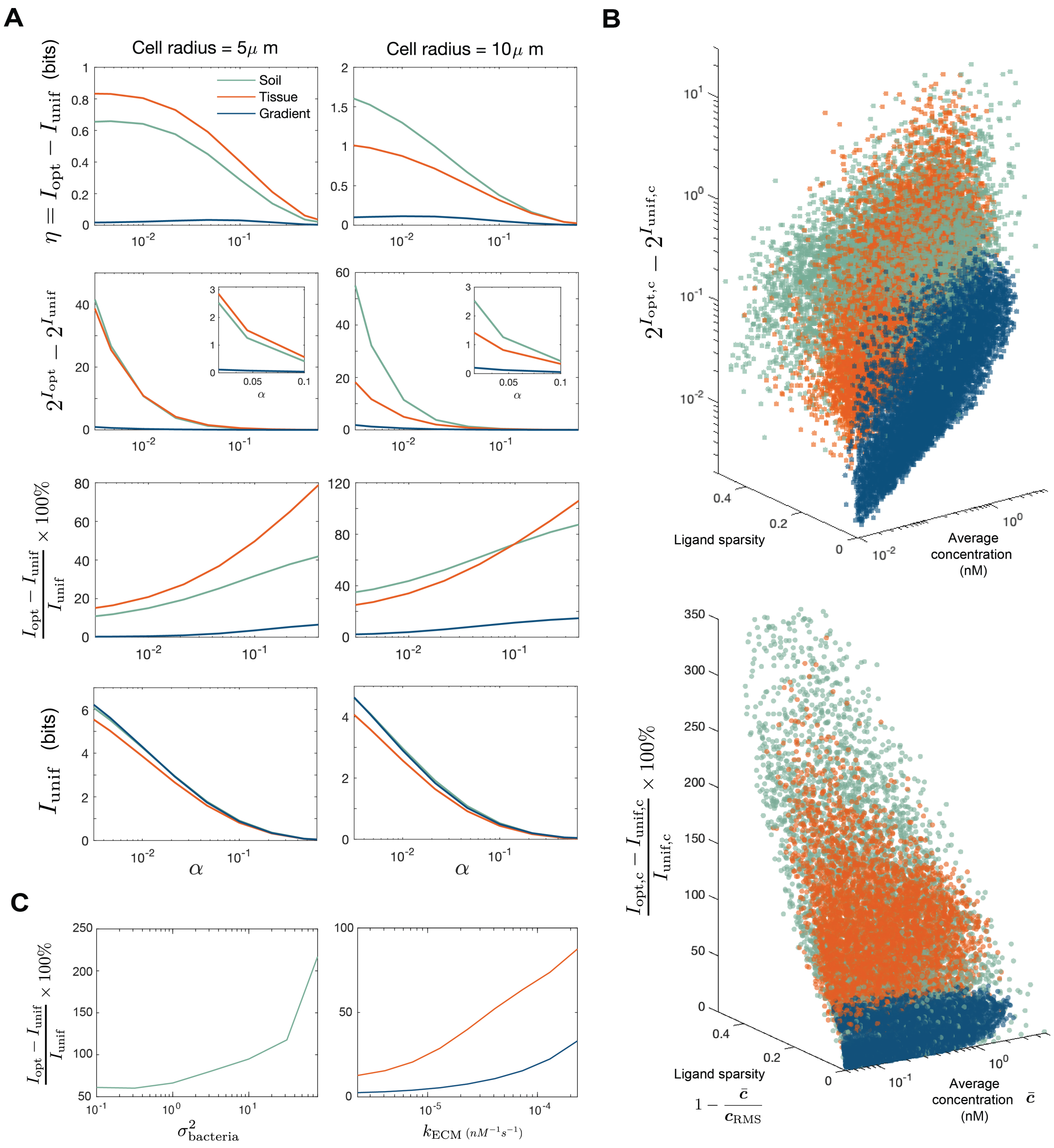


Figure S4: **Different metric assessing information gain offered by the optimal placement strategy over the uniform strategy, Related to Figure 2.** (A) versions of Figure 2C for different information metrics, where $I_{\text{opt}}$ and $I_{\text{unif}}$ is defined in Equation 5 in the main text, 1st row is absolute information gain between optimal and uniform receptors, 2nd row is absolute increase in the number of different classes to which the ligand profile can be subdivided after observing the receptor activity (with inset showing intermediate values of $\alpha$), 3rd row is relative information gain, 4th row is the average information obtained with uniform receptors; (B) versions of Figure 2E for different information metrics, $I_{\text{opt,c}} = I(\hat{\boldsymbol{c}}; \hat{\boldsymbol{a}} \mid \phi^*)$ and $I_{\text{unif,c}} = I(\hat{\boldsymbol{c}}; \hat{\boldsymbol{a}} \mid \phi^u)$; (C) versions of Figure 2D for different information measure.

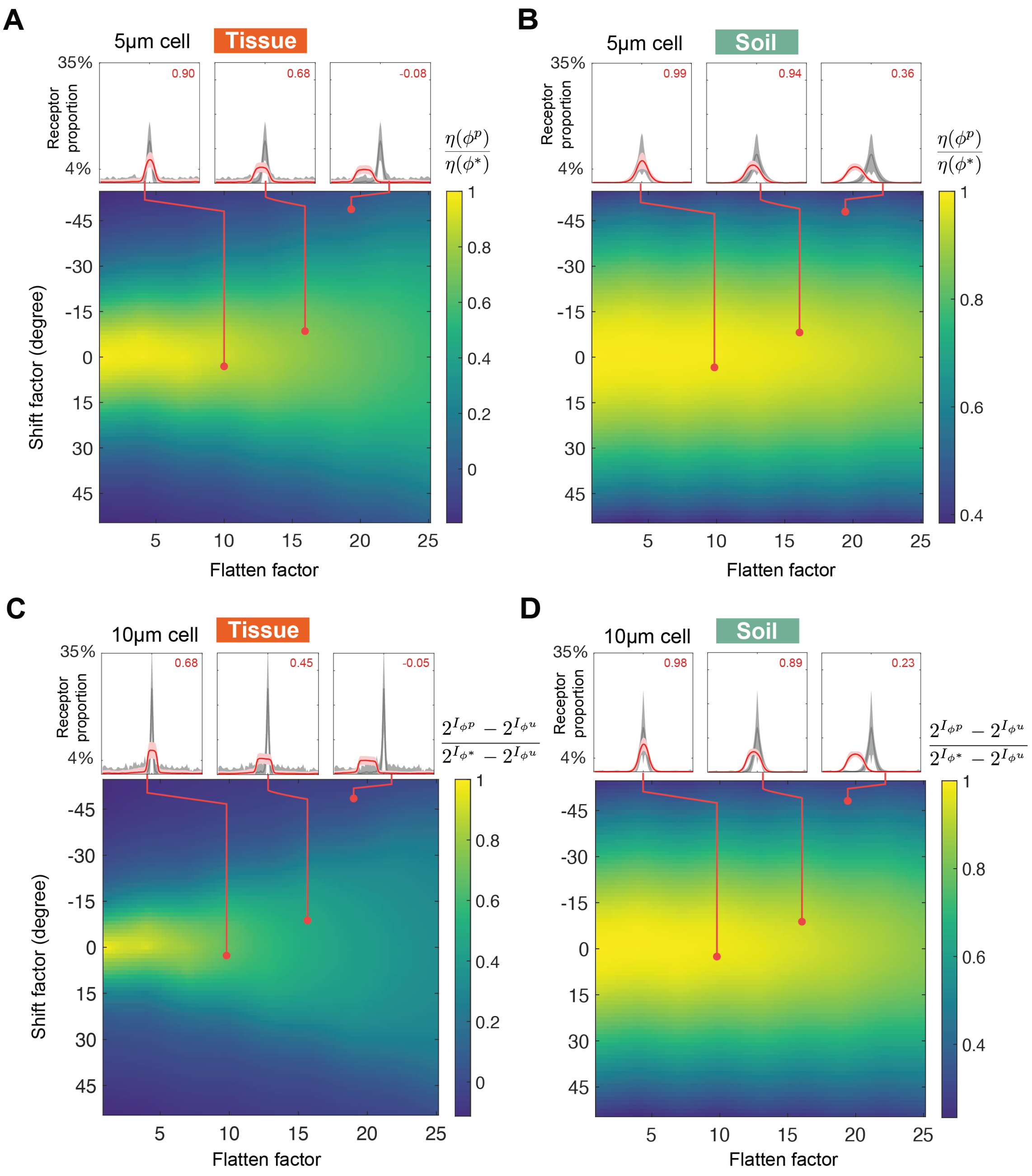


Figure S5: **Robustness of optimal efficacy to perturbation in receptor placement for other cell radius and efficacy metric, Related to Figure 3.** Colors of heat map represent ratio of perturbed efficacy $\eta(\phi^p)$ to optimal efficacy $\eta(\phi^*)$ for different combinations of shifting and flattening, computed for ligand profiles $\{\boldsymbol{c}\}$ sampled from either (A) tissue or (B) soil; call-out boxes corresponds to different sets of perturbations, showing the average of the optimal $\{\phi^*(\boldsymbol{c})\}$ (gray) and perturbed $\{\phi^p(\boldsymbol{c})\}$ (red) receptor placements, after all profile peaks were centered; (C) same as (A) but for a cell of 10 µm radius and for $\eta(\phi) := 2^{I_\phi} - 2^{I_{\phi^u}}$, denoting the increase in distinguishable input state between optimal and uniform placements, (D) similarly for soil.

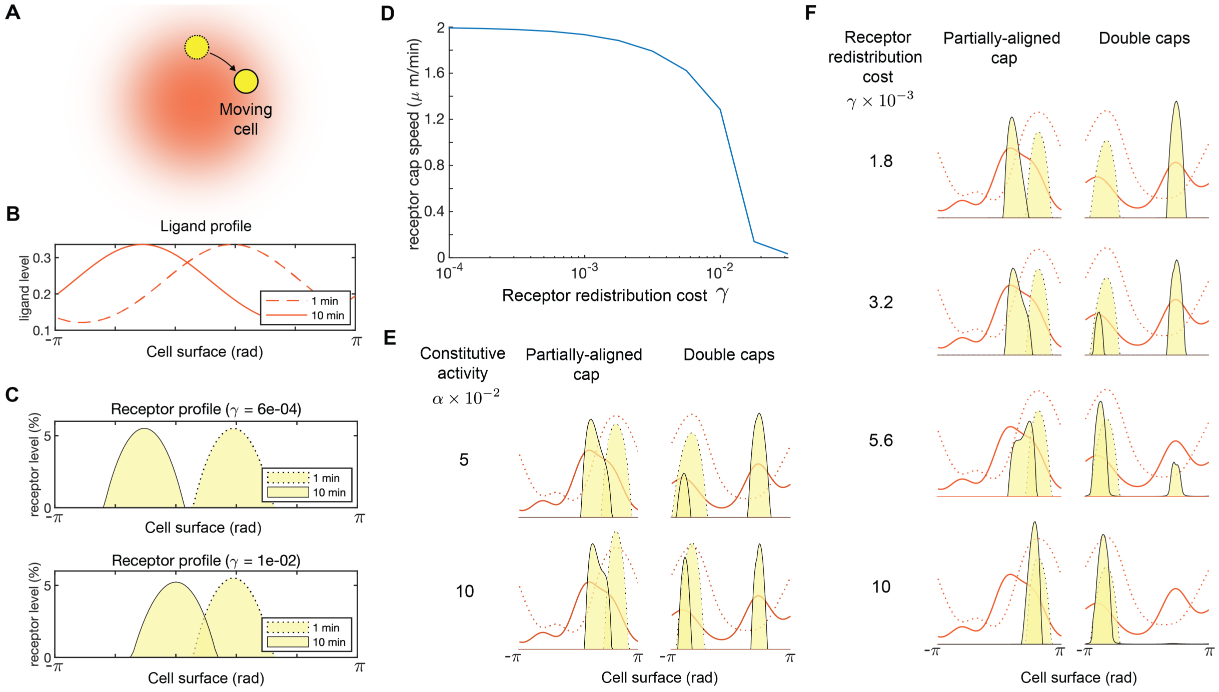


Figure S6: **Effect of redistribution cost $\gamma$ and constitutive receptor activity $\alpha$ on receptor redistribution according to the dynamic protocol, Related to Figure 4.** (A) Schematic showing a cell circling a ligand source which generates a stationary gradient (red). (B) Ligand profile experienced by the moving cell in Panel A at two different time points. (C) Optimal receptor profile computed using Equation 11 for the two time points shown in panel B , for different values of redistribution cost $\gamma$. (D) Speed of the moving receptor cap (as shown in panel C) for a wide range of $\gamma$, where speed is computed using the distance moved by the center-of-mass of the receptor distribution. (E) Different receptor redistribution dynamics for different degrees of constitutive receptor activity $\alpha$ (shown for two different pairs of ligand profiles), dotted lines represent ligand profile (red) and receptor profile (yellow shading) at one time step, while solid lines represent the ligand and receptor profile at the next time step. (F) Different receptor redistribution dynamics for different receptor redistribution cost $\gamma$, shown for the same group of ligand profiles as panel E, dotted lines represent ligand profile (red) and receptor profile (yellow shading) at one time step, while solid lines represent the ligand and receptor profile at the next time step

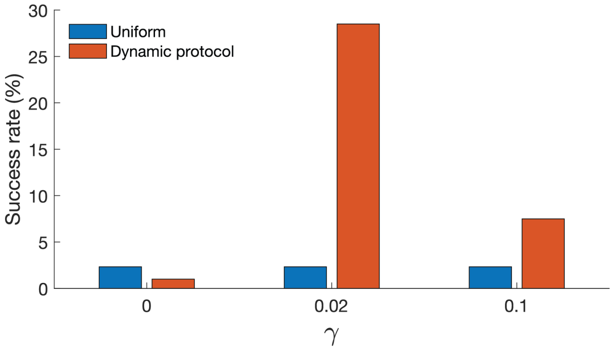


Figure S7: **Success rate of chemotactic cell navigating in simulated interstitial gradient for various values of $\gamma$ in dynamic protocol, Related to Figure 4.** Dynamic protocol of equation (S46) is solved step-wise for a cell simulated to navigate through an interstitial gradient, success rate is the proportion of simulated cells reaching gradient peak within 1 hour using the optimal + noise decoding method. $\gamma = 0$ corresponds to the case where the cost term is absent and receptors move simply to maximize mutual information. Parameter $\gamma$ determines the balance between maximizing information and minimizing receptor transport cost. Note the dynamic protocol should not be confused with the feedback scheme.

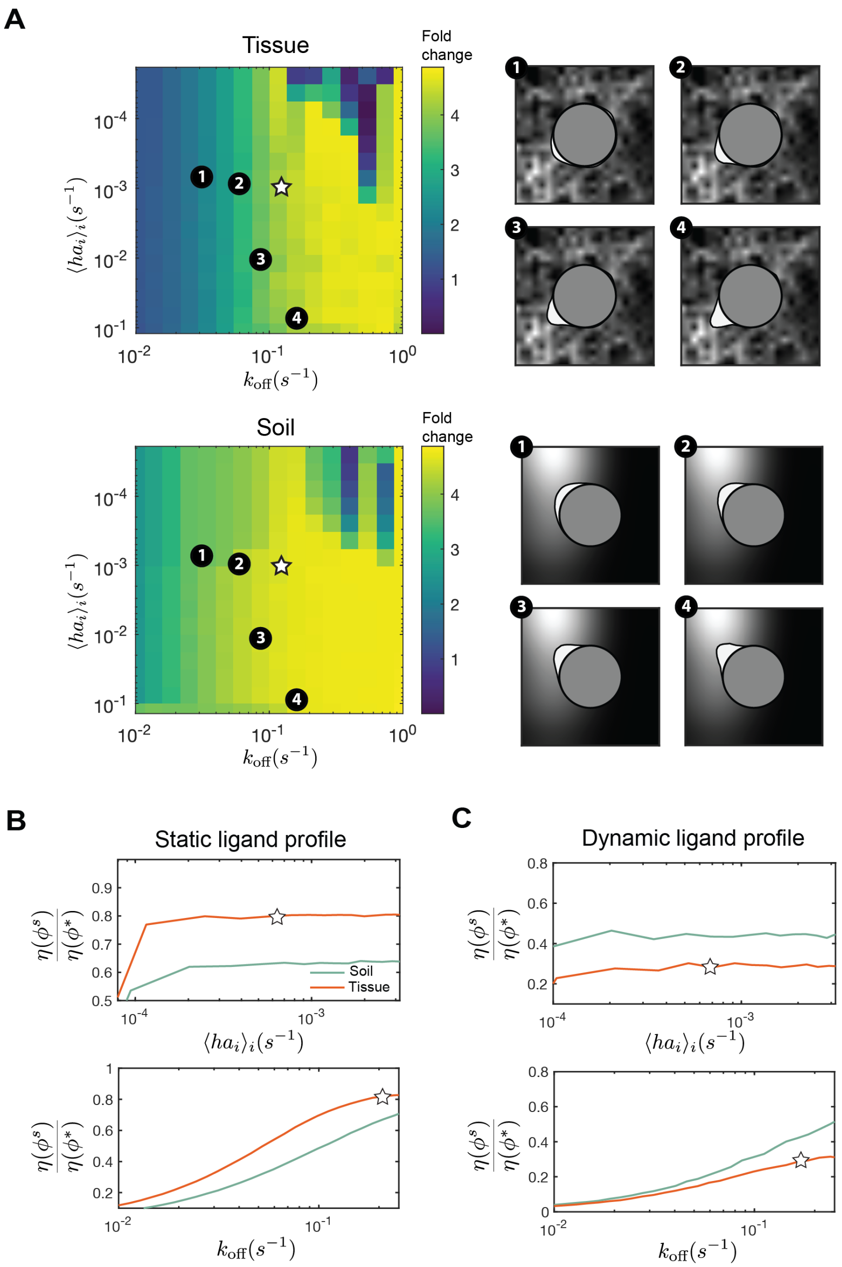


Figure S8: **Effect of rate parameter values on the effectiveness of receptor feedback scheme for information acquisition, Related to Figure 5.** (A) heat maps show the extent of receptor localization at the ligand peak for different choices of scheme parameters $k_{\mathrm{off}}$ and $h$, for a cell in tissue (top) and soil (bottom), as measured by the fold change in receptor number near the ligand peak compared to a uniform distribution of receptors; call out boxes show receptor morphology for different parameter values, star indicates default parameter values used in Figure 5. (B) ratio of scheme efficacy $\eta(\phi^s)$ to optimal efficacy $\eta(\phi^*)$ for static signals $\{\boldsymbol{c}\}$ sensed by a 5 µm cell sampled from soil and tissue (C) ratio of scheme efficacy $\eta(\phi^s)$ to optimal efficacy $\eta(\phi^*)$ for a sequence of signals $\{\boldsymbol{c}_t\}$ sampled by translating a 5 µm cell through soil and tissue environment at a speed of 2 µm min$^{-1}$.

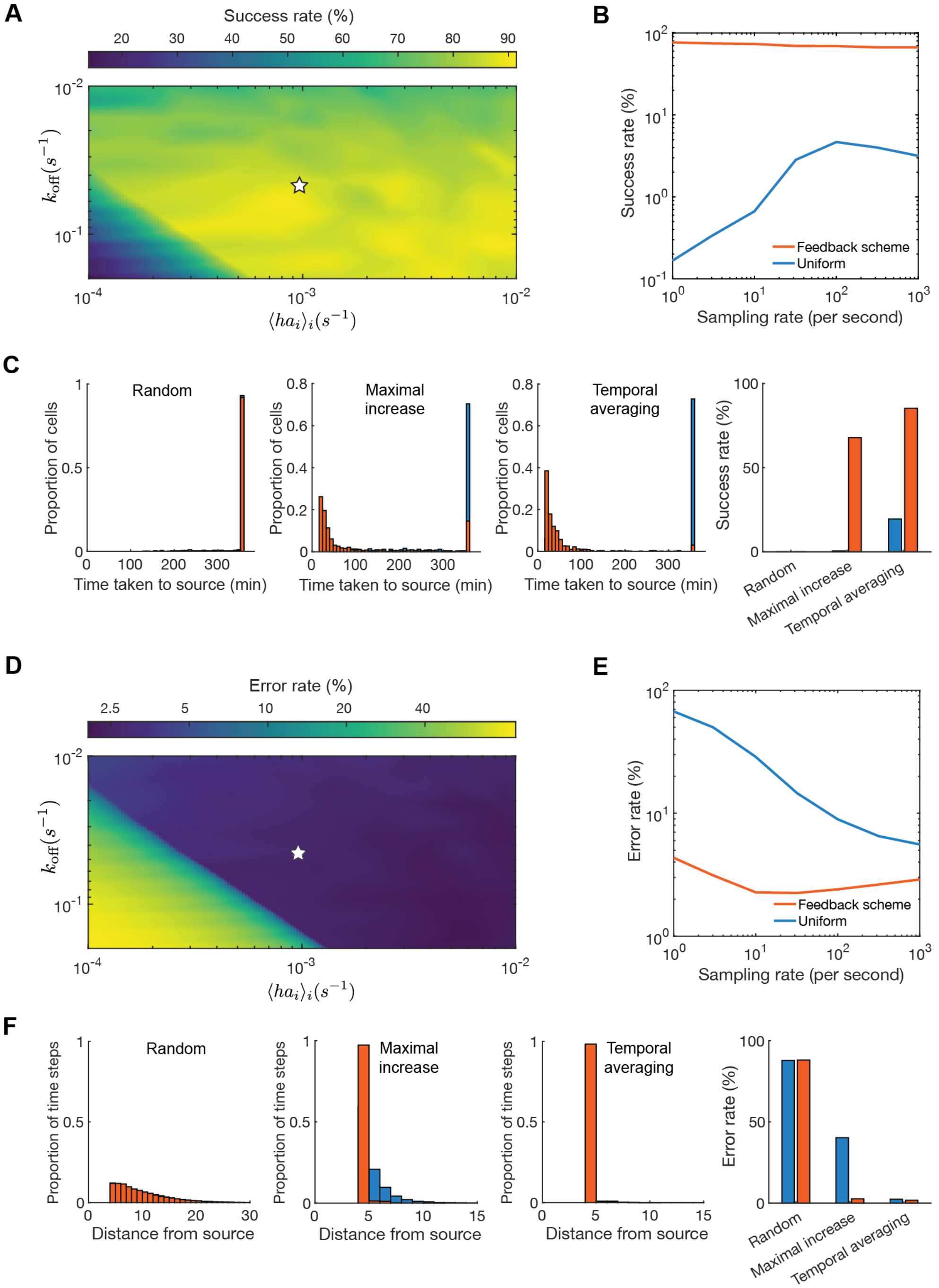


Figure S9: **Cell navigation and retention performance in simulated interstitial gradient for various feedback scheme parameters, Related to Figure 6.** (A) heatmap showing success rate (proportion of simulated cells reaching gradient peak within 1 hour) for cells using receptor feedback scheme with different values of the endocytosis rate ($k_{\text{off}}$) and average incorporation rate ($\langle ha_i \rangle_i$), white star denotes parameter values used in Figure 6 of main text. (B) success rate for cells navigating with different sampling rate, which is the number of ligand profiles sampled per second. (C) histogram and corresponding success rate quantification for cells decoding gradient using three different decoding methods, see supplemental information for detail on each method (temporal averaging is done over a 5 minute window). (D) heatmap showing error rate (proportion of simulated time steps where cell was more than $5\mu m$ away from the gradient peak) for cells using receptor feedback scheme with different values of the endocytosis rate ($k_{\text{off}}$) and average incorporation rate ($\langle ha_i \rangle_i$), white star denotes parameter values used in Figure 6 of main text. (E) error rate for cells navigating with different sampling rate, which is the number of ligand profiles sampled per second. (F) histogram and corresponding error rate quantification for cells decoding gradient using three different decoding methods, see section 7 for details

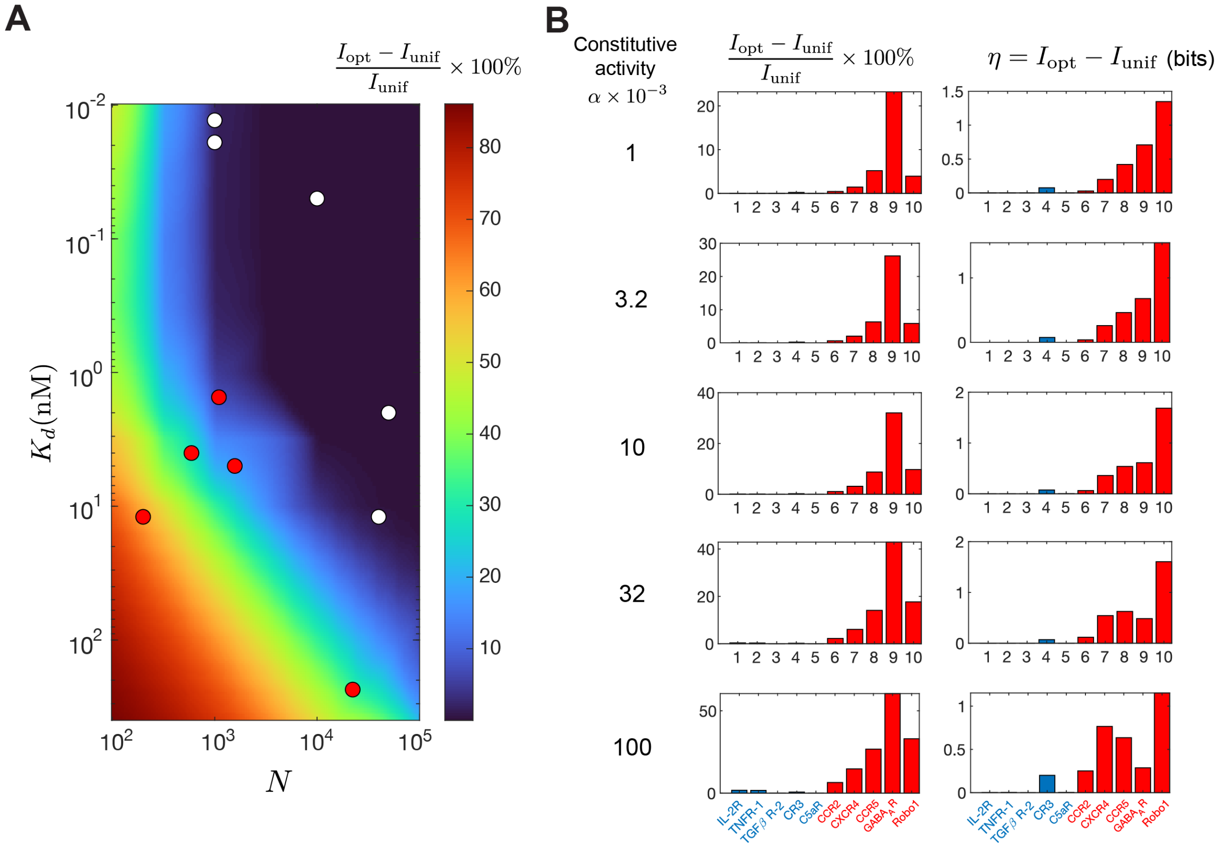


Figure S10: **Relative and absolute information gain for natural receptors and for receptors of different surface expression level, constitutive activity, and binding affinity, Related to Figure 7.** (A) relative information gain for different values of $K_d$ and $N$; values computed using the tissue environment; $\alpha = 0.1$; red dots correspond to receptors that polarize in heterogeneous environments, white dots represent receptors that are constantly uniform, see Table S4 for receptor data. (B) relative (left) and absolute information gain (right) computed for the ten natural membrane receptors shown in panel A, across different values of $\alpha$, blue bars/labels correspond to receptors that are constantly uniform (white dots in panel A), red bars/labels correspond to receptors that localize towards ligand source in concentration gradient.
